# Deciphering a marine bone degrading microbiome reveals a complex community effort

**DOI:** 10.1101/2020.05.13.093005

**Authors:** Erik Borchert, Antonio García-Moyano, Sergio Sanchez-Carrillo, Thomas G. Dahlgren, Beate M. Slaby, Gro Elin Kjæreng Bjerga, Manuel Ferrer, Sören Franzenburg, Ute Hentschel

## Abstract

The marine bone biome is a complex assemblage of macro- and microorganisms, however the enzymatic repertoire to access bone-derived nutrients remains unknown. The bone matrix is a composite material made up mainly of organic collagen and inorganic hydroxyapatite. We conducted field experiments to study microbial assemblages that can use organic bone components as nutrient source. Bovine and turkey bones were deposited at 69 m depth in a Norwegian fjord (Byfjorden, Bergen). Metagenomic sequence analysis was used to assess the functional potential of microbial assemblages from bone surface and the bone eating worm *Osedax mucofloris*, which is a frequent colonizer of whale falls and known to degrade bone. The bone microbiome displayed a surprising taxonomic diversity revealed by the examination of 59 high-quality metagenome assembled genomes from at least 23 bacterial families. Over 700 genes encoding enzymes from twelve relevant enzymatic families pertaining to collagenases, peptidases, glycosidases putatively involved in bone degradation were identified. Metagenome assembled genomes (MAGs) of the class Bacteroidia contained the most diverse gene repertoires. We postulate that demineralization of inorganic bone components is achieved by a timely succession of a closed sulfur biogeochemical cycle between sulfur-oxidizing and sulfur-reducing bacteria, causing a drop in pH and subsequent enzymatic processing of organic components in the bone surface communities. An unusually large and novel collagen utilization gene cluster was retrieved from one genome belonging to the gammaproteobacterial genus *Colwellia.*

**Importance:** Bones are an underexploited, yet potentially profitable feedstock for biotechnological advances and value chains, due to the sheer amounts of residues produced by the modern meat and poultry processing industry. In this metagenomic study we decipher the microbial pathways and enzymes that we postulate to be involved in bone degradation marine environment. We herein demonstrate the interplay between different bacterial community members, each supplying different enzymatic functions with the potential to cover an array of reactions relating to the degradation of bone matrix components. We identify and describe a novel gene cluster for collagen utilization, which is a key function in this unique environment. We propose that the interplay between the different microbial taxa is necessary to achieve the complex task of bone degradation in the marine environment.

## Introduction

The marine environment is a treasure trove for novel microbial assemblages and organic catalysts (enzymes) (1–3). The oceans cover approximately 70% of the Earth surface with an estimated volume of about 2 × 10^18^ m^3^ and due to its incredible environmental variability (e.g. temperature, pressure, salinity, light availability), it has sparked the evolution of an unprecedented range of different microbes and hence enzymatic activities (4–6). Genome sequencing of individual microbial isolates of complex communities has allowed us to get a glimpse of their diversity and their potential functions in multiple environmental contexts. The lack of cultivable microbes has further driven the development of functional and sequence-driven metagenomic analyses, and enabled us to decipher complex interactions in entire microbial consortia (7–9).

The deep-sea was for a long time seen as an almost lifeless environment, as no one could imagine life to be possible under conditions vastly different from shallower ocean waters in respect to nutrient and energy resources. Nowadays we know that even the deep-sea is teeming with life; hydrothermal vents, sponge grounds and coral gardens are recognized as examples of unique and complex habitats (10–12). Nonetheless, the deep-sea is also a harsh environment with limited nutrient sources. In this respect sudden events like a whale fall, create a locally defined but significant nutrition source for deep-sea life, that can last for years or even decades (13). These whale carcasses are rapidly stripped of their soft tissue by scavengers (i.e., hagfish, sleeper sharks, rat tail fish, crabs), but the energy-rich bones and cartilage remain as a recalcitrant nutrient source. More than 15 years ago *Osedax* was described, a genus of bone-eating annelid worms (14), and has since then been investigated for its diversity, ecology and how it accesses the organic compounds of whale bones (14–17). These worms are gutless and rather bore cavities into bones and develop a root tissue in these cavities for food intake. Furthermore, this evolutionary novel and specialized organ was shown to harbor endosymbionts, typically affiliated to Oceanospirillales (14, 18–21). *Osedax* species are known for their ability to acidify their environment via elevated expression levels of vacuolar-H^+^-ATPase (VHA) specifically in their root tissue and of carbonic anhydrase throughout their body, to dissolve hydroxyapatite and access collagen and lipids from the bone matrix (16). Miyamoto *et al.* found a high number of matrix metalloproteinases in the genome of *Osedax japonicus* compared to other invertebrates, potentially assisting in digestion of collagen and other proteins derived from bones (15). The species can thus be regarded as a member of the bone biome and an important facilitator in this degradation process. In the northern North Atlantic *Osedax mucofloris* was described in 2005 and has been shown to consistently colonize bone material on the sea floor below a depth of 30 m (22, 23).

Bone is a recalcitrant and heterogeneous composite material made of a mineral phase, an organic phase and water. Hydroxyapatite crystals in the mineral phase contribute to the structural strength in bones. The organic phase includes proteins, such as collagen and structural glycoproteins (e.g. proteins decorated with sugars such as mannose, galactose, glucosamine, galactosamine, N-acetylglucosamine, N-acetylgalactosamine, rhamnose, sialic acid and fucose), lipids and cholesterol composed of various triglycerides (24–26). Up to 90% of the protein content in mature bone is made of type I collagen, a triple helical molecule rich in glycine, hydroxyproline and proline that assembles into fibrils with a high number of hydrogen bonds, hydrophobic interactions and covalent cross-linking, which together confer high structural stability to collagen fibrils (27). Due to this structural and chemical complexity, it is expected that the degradation of the recalcitrant bone matrix will require a synergistic multi-enzyme system and require a microbial community effort. Similar multi-enzyme systems are well described for example for the degradation of lignocellulose, another important organic polymer (28, 29). This system will likely include essential enzymes in the breakdown of the organic matrix, namely collagenases that break the peptide bonds in collagen and other proteases/peptidases (endo- and exopeptidases) that attack the glycoproteins. Furthermore, neuraminidases (sialidases), α-mannosidases, α-/β-galactosidases, α-fucosidase, α-rhamnosidase and α/β-N-acetylhexosaminidase (glucose and galactose-like), all glycoside hydrolase enzymes are likely involved in cleavage of glycosidic linkages. Finally, in the digestion of the cholesterol-containing marrow, cholesterol oxidases may be involved.

To date only a few studies have been published that focus on microbial communities to understand the necessary complex interactions in bone degradation, mainly relying on 16S rRNA gene sequencing data (30–33) and one metagenomic study of a whale fall (34). We here provide a first comprehensive overview and identify putative key functions involved in bone degradation of the marine bone microbiome retrieved from deployed bone material, including microbial communities from the gutless worm *Osedax mucofloris* and free-living microbial assemblages developing on the bone surface.

## Results

### Recovery of artificially deployed bone for bone microbiome metagenomic analysis

Turkey and bovine bones were deployed at 69 m depths in Byfjorden, a fjord outside Bergen, Norway. After nine months of incubation, underwater images taken by a remotely operated vehicle (ROV) showed microbial colonization of the bone surfaces (supplementary figure S1A). *Osedax mucofloris* worms were observed, especially on the joints, and in some cases forming small colonies of several individuals inside a single cavity under the external bone tissue. Although not subject of this study, a larger diversity of invertebrate fauna including *Ophryotrocha, Vigtorniella* and *Capitella* worms were also observed. Dense microbial mats developed asymmetrically with preference for the joint adjacent sections (epiphysis), which also appeared in aquaria settings (Supplementary figure S1B).

Two sets of samples, an *Osedax-*associated bone microbiome (OB), and a bone surface-associated biofilm (BB) were collected, each consisting of four individual metagenomes (Table 1). The co-assemblies of each sample set comprised of >300 000 contigs and 342.7 Mb for the OB-metagenomes and >1 000 000 contigs and 1.22 Gb for the BB-metagenomes respectively, considering only contigs >500 bp (Supplementary table S1). The individual metagenomes were profiled separately with Kaiju (35), a database aided metagenomic read taxonomy classifier. The OB-metagenomes comprised of 45-76% bacterial, 15-34% eukaryotic and 0.4-0.5% archaeal reads and the BB-metagenomes are made up of 92-95% bacterial, 2-4% eukaryotic and 0.4-0.7% archaeal reads (Supplementary table S1).

**Table 1:**
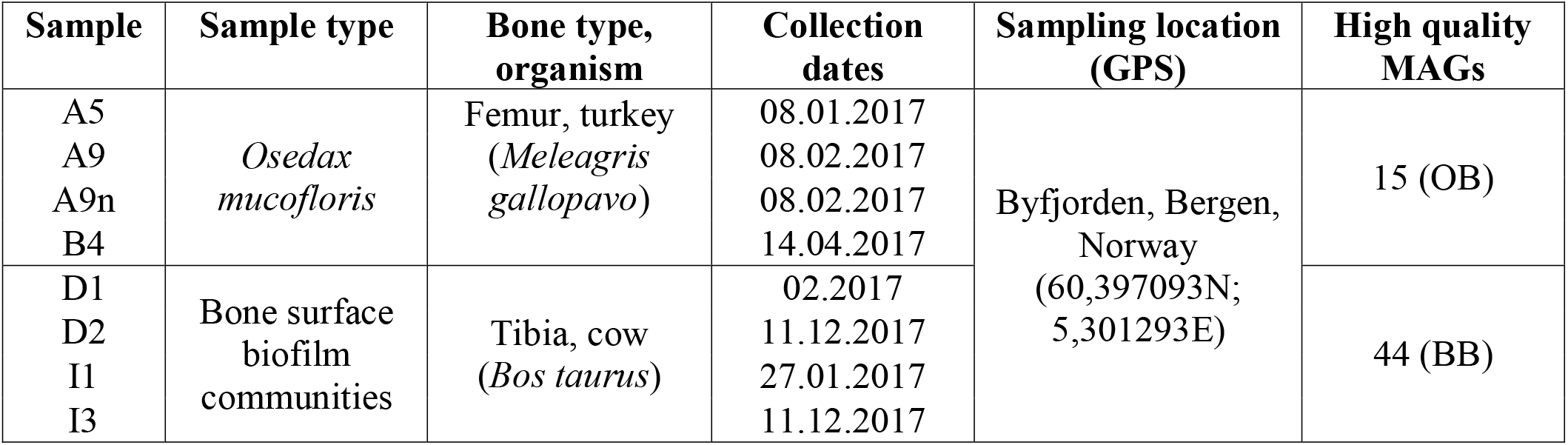
Metagenome sampling information and number of retrieved metagenome assembled genomes (MAG).

| Sample | Sample type | Bone type, organism | Collection dates | Sampling location (GPS) | High quality MAGs |
| --- | --- | --- | --- | --- | --- |
| A5<br>A9<br>A9n<br>B4 | *Osedax mucofloris* | Femur, turkey<br>(*Meleagris gallopavo*) | 08.01.2017<br>08.02.2017<br>08.02.2017<br>14.04.2017 | Byfjorden, Bergen, Norway<br>(60,397093N;<br>5,301293E) | 15 (OB) |
| D1<br>D2<br>I1<br>I3 | Bone surface biofilm communities | Tibia, cow<br>(*Bos taurus*) | 02.2017<br>11.12.2017<br>27.01.2017<br>11.12.2017 |  | 44 (BB) |

### Metagenome assembled genomes (MAGs) from the marine bone microbiome display taxonomic diversity and novelty

59 high-quality MAGs (>90% completion and <10% redundancy) were extracted from the coassembled metagenomes (Supplementary Table S3 for MAG sequence statistics). The MAGs span 11 phyla, 14 classes, 19 orders and at least 23 families. About 63% of the MAGs (37/59) are taxonomically novel as determined by their relative evolutionary divergence (RED) (36) to their closest common ancestor (Supplementary table S2). One MAG could be only identified up to phylum level, seven to class level, seven to order level, 18 up to family level and four up to genus level. The taxonomy of most MAGs was fully resolved based on 120 marker genes. The three best represented phyla were Proteobacteria (22 MAGs), Campylobacterota (14 MAGs) and Bacteroidota (8 MAGs). However, the percental distribution of the most abundant classes differs between the two metagenome sets. The OB-MAGs were dominated by the classes Gammaproteobacteria (27%), Campylobacteria (27%) and Alphaproteobacteria (20%), while the BB-MAGs were mainly affiliated with Gammaproteobacteria (30%), Campylobacteria (23%) and Bacteroidia (16%).

### Sulfur cycling in the marine bone microbiome

All MAGs were investigated using multigenomic entropy based score pipeline (MEBS) (37) for their ability to utilize sulfur as energy source via the abundance of selected marker genes for related pathways (Figure 2). Sulfur cycling is of relevance to bone degradation due to the generation of free protons by sulfur and sulfur compound oxidation processes (thiotrophy), which leads to an acidification.

MAGs affiliated to the order Campylobacterales encode almost complete Sox (*sox*XYZABCD) and Sor enzyme systems for sulfur oxidation. Furthermore, Gammaproteobacteria affiliated to Beggiatoaceae and other unclassified Gammaproteobacteria, are all potentially capable of thiotrophy via utilization of reduced sulfur compounds as electron donors (flavocytochrome c sulfide dehydrogenase *fcc*B) and adenosine-5’-phospho-sulfate reductase (*apr*AB)) and partial predicted Sox sulfur/thiosulfate oxidation pathway. MAGs identified as Desulfuromonadia, Desulfobacteria, Desulfobulbia and Desulfovibrionia possess marker genes for sulfur reduction *(qmoABC, hydACD, sre*ABC) and lack Sox and Sor pathway genes, all of which belong to the new proposed phylum of Deltaproteobacteria (38). In all Gammaproteobacteria (except BB32), Desulfobacteria, Desulfobulbia and Desulfovibrionia, genes for dissimilatory sulfite reductase (*dsr*ABC) are present. Müller *et al.* (2015) described that gammaproteobacterial *dsr*AB-type genes are commonly involved in oxidative reactions, whereas *dsr*AB in Desulfobacterota are reductive type *dsr*AB (39). All MAGs contain at least partial pathways for dissimilatory tetrathionate reduction (*ttr*ABC), thiosulfate disproportionation (*phs*ABC and rhodanase), dimethylsulphide (DMS) degradation (*ddh*ABC) and contain also genes for sulfoacetaldehyde degradation *(isfD,* xsc and *saf*D). In addition, the phenotypic trait of H_2_S production was identified in 10 MAGs (Traitar analysis (40)), two of which were Marinifiliaceae, two Krumholzibacteria, two *Sulfurospirillum,* one Spirochaetaceae, two Desulfobacteraceae, and one *Pseudodesulfovibrio*.

The anticipated thiotrophy has the potential to contribute massively to the acidification of the environment via the oxidation of reduced sulfur compounds leading to production of sulfuric acid (41). This requires a close interaction between sulfur-reducing bacteria (SRB) producing hydrogen sulfide and sulfur-oxidizing bacteria (SOB) utilizing the hydrogen sulfide, while releasing protons (Figure 3). The Traitar analysis identified 10 MAGs potentially able to produce hydrogen sulfide, including known SRB like Desulfobacteraceae, *Pseudodesulfovibrio,* and others like *Sulfurospirillum* (42–44). The bone microbiome is especially enriched in taxa containing known SOB, like the large filamentous bacteria Beggiatoales (5 MAGs) (45) and Campylobacterales (10 MAGs) (46). Furthermore, one MAG identified as Desulfobulbaceae was found in the bone associated metagenomes. Member of this group are known to be able to perform sulfur oxidation and sulfur reduction (47, 48).

### Acidification by carbonic anhydrases

Carbonic anhydrases were identified in 51 of 59 MAGs. Nineteen out of 94 carbonic anhydrases contained a signal peptide for extracellular export (16 MAGs). 15 were predicted to contain a Sec signal peptide (SPI) and four to encode lipoprotein (SPII) signal peptides. Four out of five Beggiatoales MAGs were predicted to contain carbonic anhydrases with a SPI signal peptide (BB2, BB3 and BB20) or a SPII signal peptide (BB16). The remaining three SPII signal peptides were found in carbonic anhydrases from Campylobacterales (BB8, BB10 and BB11) Interestingly BB8 contains at least three carbonic anhydrases, one with a SPI, one with a SPII and one where a signal peptide was not predicted. Three SPI signal peptides were found in carbonic anhydrases from unclassified γ-proteobacteria (BB4, BB34 and BB36). The remaining SPI including carbonic anhydrases were found in five Campylobacterales MAGs (BB14, BB26 (three carbonic anhydrases and two containing SPI signal peptides), BB30, BB41 and OB7) and one Desulfobulbaceae MAG (BB13). Based on phylogenetic relationship to known carbonic anhydrases described in Capasso *et al.,* 2015 (49), 18 out of 19 carbonic anhydrases belong to the α-carbonic anhydrase family and one to the β-family, with no γ-family carbonic anhydrases found (Supplementary figure S4).

### Gelatin hydrolysis

With respect to microbial bone degradation, the phenotypic feature of gelatin hydrolysis was analyzed using the Traitar software (40), which provides genome-informed phenotype predictions. 22 MAGs showed capacity of gelatin hydrolysis (19 MAGs in the bone surface community (BB) and three in the *Osedax* associated communities (OB)). With gelatin being a primarily bone collagen derived compound, we consider gelatin hydrolysis a key trait for the microbial community studied herein. All eight Bacteroidia affiliated MAGs possess the gelatin hydrolysis trait, seven Gammaproteobacteria MAGs, one tentative Planctomycetota MAG, one Spirochaetia MAG, two Krumholzibacteria MAGs, one Thiovulaceae, one Geopsychrobacteraceae and one Fermentibacteria MAG (Figure 3).

### Enzymes involved in bone degradation

Based on the structure and composition of mature vertebrate bone tissue, we hypothesized that 12 different COGs and peptidase/collagenase families were relevant for the enzymatic attack of the bone organic matrix. This “bone-degradome” comprised the peptidase families S1 (COG0265), S8/S53 (and Pfam00082), U32 (COG0826) and M9 collagenase (including Pfam01752), mannosidases (COG0383), sialidases (COG4409), glucuronidases (COG3250), glucosaminidases (COG1472), galactosaminidases (COG0673), α-galactosidases (Pfam 16499), cholesterol oxidases (COG2303) and fucosidases (COG3669) (choice of enzymes is justified in the Materials and Methods section). We constructed HMM profiles that were used to screen the abundance of each enzyme family in all MAGs (Figure 4). In total 722 enzymes belonging to the 12 investigated enzyme families were identified in the 59 MAGs. The glycosidase families of mannosidases, galactoaminidases, glucosaminidases and the peptidase families S1, S8/S53 and U32 were widespread in the MAGs (Figure 4). M9 collagenases and α-galactosidases (Pfam 16499) were only found in three MAGs. The M9 collagenases where solely found in Enterobacterales (BB5, BB44 and OB 12). Pfam16499 was only identified in Bacteroidales (BB22, BB24 and OB13). The most abundant group of enzymes were the S1 peptidases (141 hits), followed by galactosaminidases (COG0673) (116 hits) and U32 peptidases (99 hits) (Figure 4), constituting 20%, 16% and 14% of all identified bone degrading enzymes, respectively. In general, Bacteroidales (BB17, BB22, BB24, BB29, BB42 and OB13) displayed the most diverse set of enzyme families related to bone degradation, as they contained genomic evidence of all enzymes besides M9 collagenases. MAGs belonging to the orders Desulfuromonadia, Desulfobulbia, Desulfobacteria, Desulfovibrio, Campylobacteria (all of them driving the sulfur biogeochemical cycle), as well as some undefined α-proteobacteria and γ-proteobacteria appear to have no or few mannosidases (COG0383), glucuronidases (COG3250), fucosidases (COG3669), sialidases (COG4409) and α-galactosidases (Pfam16499).

### Collagen degradation

We investigated the genomic context of each M9 collagenase for potential links to metabolic pathways, such as proline utilization (Supplementary figure S4). *Colwellia* MAG BB5 possessed an approximately 21 kbp-long gene cluster presumably devoted to collagen utilization, which is unique in the dataset and in the public databases. The functional cluster spans at least 15 different genes (Figure 5A), featuring a secreted Zn-dependent M9 collagenase, a secreted peptidyl-prolyl cis-trans isomerase (cyclophilin-type PPIase), a secreted unknown protein and an unknown Zn/Fe chelating domain-containing protein. Additionally, one putative transporter (MFS family), a TonB-dependent receptor and several genes involved in the catabolism of proline and hydroxyproline, e.g. prolyl-aminopeptidase YpdF, intracellular peptidyl-prolyl cistrans isomerase (rotamase), pyrroline reductase, hydroxyproline dipeptidase, 4-hydroxyproline epimerase and others. Moreover, genes involved in transcription regulation such as PutR and the stringent starvation protein A and B were identified in the same putative gene cluster.

To explore the conservation of this gene cluster, we retrieved fourteen representative *Colwellia* genomes of marine origin from the NCBI repository (Supplementary table S5) (50–63). To minimize methodological bias, the nucleotide sequences of these genomes were likewise annotated with RAST (rapid annotation using subsystem technology) and screened for M9 collagenase using the previously established HMM profile. 22 annotated M9 collagenases were identified in seven out of fourteen genomes. In the genomes of *Colwellia piezophila* ATCC BAA-637 (57) and *Colwellia psychrerythraea* GAB14E (59) a gene cluster comparable to the one in MAG BB5 was identified (Figure 6) and found to be largely conserved between the three species. The conserved core constitutes of the M9 collagenase, a D-hydroxyproline dehydrogenase, an epimerase, a secreted cyclophilin-type peptidyl-prolyl isomerase, a ketoglutaric acid dehydrogenase, a MFS transporter, an aminopeptidase, a bifunctional 1-pyrroline-5-carboxylate reductase/ornithine cyclodeaminase and a hydroxyproline dipeptidase. BB5 additionally contains several other relevant genes, such as a PutR regulator, stringent starvation proteins A and B, a TonB dependent receptor, Zn/Fe binding domain protein, 1-pyrroline-4-hydroxy-2-carboxylate deaminase and an intracellular peptidyl-prolyl cis-trans isomerase (rotamase, PpiD).

### Specificity of the bone microbiome

To investigate the potential specificity of the bone microbiome in respect to its taxonomic composition and function the generated MAGs were compared to 832 seawater MAGs generated from the Tara Ocean datasets, comprising 93 metagenomes from various locations (64). The bone microbiome was rebinned to retrieve all MAGs with >70% completion to match the Tara Ocean threshold for high-quality MAGs, resulting in 86 MAGs. The taxonomic class abundances were compared between the two datasets and Gammaproteobacteria and Bacteroidia appeared in similar abundances (bone microbiome 23.26% and 15.12% and Tara Oceans 23.56% and 13.34% respectively), but other than those high taxonomic ranks, the datasets were different. The bone microbiome is dominated by Campylobacteria, accounting for 26.74% of the MAGs, whereas only 0.12% Campylobacteria MAGs are found in the Tara Oceans dataset. In contrast Alphaproteobacteria are better represented in the Tara Oceans dataset, than in the bone microbiome, 21.88% to 5.81% respectively. The bone microbiome furthermore also contains a number of bacterial classes that are represented at low levels (1-5% of the MAGs), which are either not represented in the Tara Oceans dataset or only at miniscule levels as low as 0.1-0.5% (Supplementary figure S4A). The functional repertoire was compared via screening the Tara Oceans dataset with the previously generated HMM profiles for enzymes potentially involved in bone degradation and calculated as ratio of enzymes per MAG. The ratios for 9 out of 12 profiled enzyme families were higher in the bone microbiome, ratios for α-N-acetylgalactosaminidases (COG0383) and cholesterol oxidases (COG2303) were higher in the Tara Oceans dataset and the ratio for S1 peptidases was equal between the two MAG sets (Supplemetary figure S4B).

## Discussion

In this study, 59 high quality MAGs were reconstructed from microbes colonizing bone surfaces and from symbionts of the bone-eating worm *Osedax mucofloris.* Metabolic reconstruction revealed a complex, diverse and specialized community. Our MAGs span at least 23 bacterial families and uncover a large potential for taxonomic novelty (over 50% according to genome-based taxonomy) from species up to class level in the bone microbiome. Interestingly, only genomes of gram-negative bacteria were reconstructed and despite gram-positive bacteria being widespread in the marine environment, they make up only minor portions of the metagenomes (4-5% of the reads affiliated to Actinobacteria and Firmicutes in the *Osedax* metagenomes and 5-9% in the Biofilm metagenomes respectively) (65). This is remarkable since they are known to carry out potentially relevant metabolic processes (thiotrophy, sulfidogenesis) (66, 67), are capable of dealing with low pH conditions which are likely encountered during bone dissolution (68), and they possess high capacity for the secretion of hydrolytic enzymes (69). Despite this underrepresentation of gram-positive taxa, this study reveals the existence of a specialized bone-degrading microbiome in the marine environment and starts to explore the enzymatic activities involved in the complete demineralization of bone material. The bone microbiome is different from seawater communities and from other specialized habitats (recolonized volcanic eruption site) in its microbial composition as well as functional makeup (Supplementary figure 4).

### The role of *Osedax* endosymbionts in bone utilization

Two distinct bacterial endosymbiont genomes belonging to the order Oceanospirillales have previously been sequenced, but their role in bone degradation in the marine environment remained unclear (18). The bacterial fraction of the herein sequenced *Osedax mucofloris* metagenome is made up of 7% to 22% Oceanospirillales affiliated reads, whereas the bone surface metagenome only contains 1% to 4% reads of this order according to the performed Kaiju analysis. This relative difference confirms that the methodological approach to minimize cross-contamination was successful and that the OB-MAGs affiliated to Oceanospirilalles likely represent the symbiotic community of *Osedax mucofloris* worms. Two MAGs belonging to the Oceanospirillales were identified in the *Osedax*-associated metagenome, belonging to the genera *Neptunomonas* (OB1) and *Amphritea* (OB2). Both genera are known to have an aerobic organotrophic metabolism and also able to thrive as free-living bacteria (70). In fact, the scarce representation of Oceanospirillales in the bone surface has been reported before (30, 31) and contrasts with their character as common dwellers of the marine environment (71, 72). Although we cannot rule out that Oceanospirillales preferentially colonize bone surfaces in earlier or later stages, their preference for a symbiont life supports the notion of a casual and facultative association to *Osedax* worms, triggered by the common benefit from a sudden nutrient bonanza as previously hypothesized (18, 21).

### The degradative functions within the bone microbiome

#### Acidification via a closed sulfur biogeochemical cycle

Free-living microbial communities, must deal with similar challenges as the *Osedax* holobiont to access the nutrient-rich, collagen-made organic bone matrix and eventually the lipid-rich bone marrow by dissolving the hydroxyapatite. The association in large specialized microbial consortia may be a beneficial strategy for achieving this task. We hypothesize that sulfur-driven geomicrobiology (sulfate/thiosulfate/tetrathionate reduction and sulfide/sulfur/thiosulfate oxidation) is the major responsible factor for bone dissolution in the marine environment by free-living bacterial communities. Campylobacterales are one of the most abundant bacterial orders in the herein investigated metagenomes, in both the *Osedax-*associated metagenomes (OB) and the bone surface biofilms (BB). Campylobacterales represent the most abundant group in terms of absolute read number, although it is the second largest taxon with reconstructed MAGs (Figure 1). Members of the Campylobacterales have previously been found to be associated with *Osedax,* albeit not as endosymbionts (21). The majority of retrieved Campylobacterales MAGs (14 in total) belong to different families of aerobic and facultative anaerobic (nitrate, manganese) sulfur-oxidizing bacteria (73, 74) (Thiovulaceae, Sulfurovaceae and Arcobacteraceae). Other aerobic/facultatively anaerobic (nitrate) sulfur oxidizing bacteria are also well represented in the order Beggiatoales (Gammaproteobacteria, 5 MAGs). *Beggiatoa-like* bacterial mats are commonly associated with whale falls (75) indicating a potential indifference regarding the bone type they dwell on. Sulfide oxidation produces elemental sulfur or sulfate (41, 45) while releasing protons and thereby causing a drop in pH. This acidification mechanism has been linked to bone demineralization. The dissolution of the hydroxyapatite mineral exposes the organic matrix to enzymatic degradation (30, 31). Besides thiotrophy, that seems to be a major acid-producing mechanism in the microbial community, other mechanisms might also contribute significantly. In this respect, a number of carbonic anhydrases (CA) were annotated, normally housekeeping genes involved in internal pH homeostasis and other processes (76), but known to play a role in environmental acidification by *Osedax* (15). Here, the CAs were found to contain signal peptides for extracellular export (19 out of 94) and therefore could also be involved in acidification. Interestingly, 18 out of 19 identified and potentially secreted CAs belong to the α-CA family and only one member of the β-CA family was found (Supplementary figure S4). The α-CA family is only found in gram negative bacteria, which is also the case here, and it is evolutionarily the youngest of the three bacterial CA families (49).

**Figure 1:**
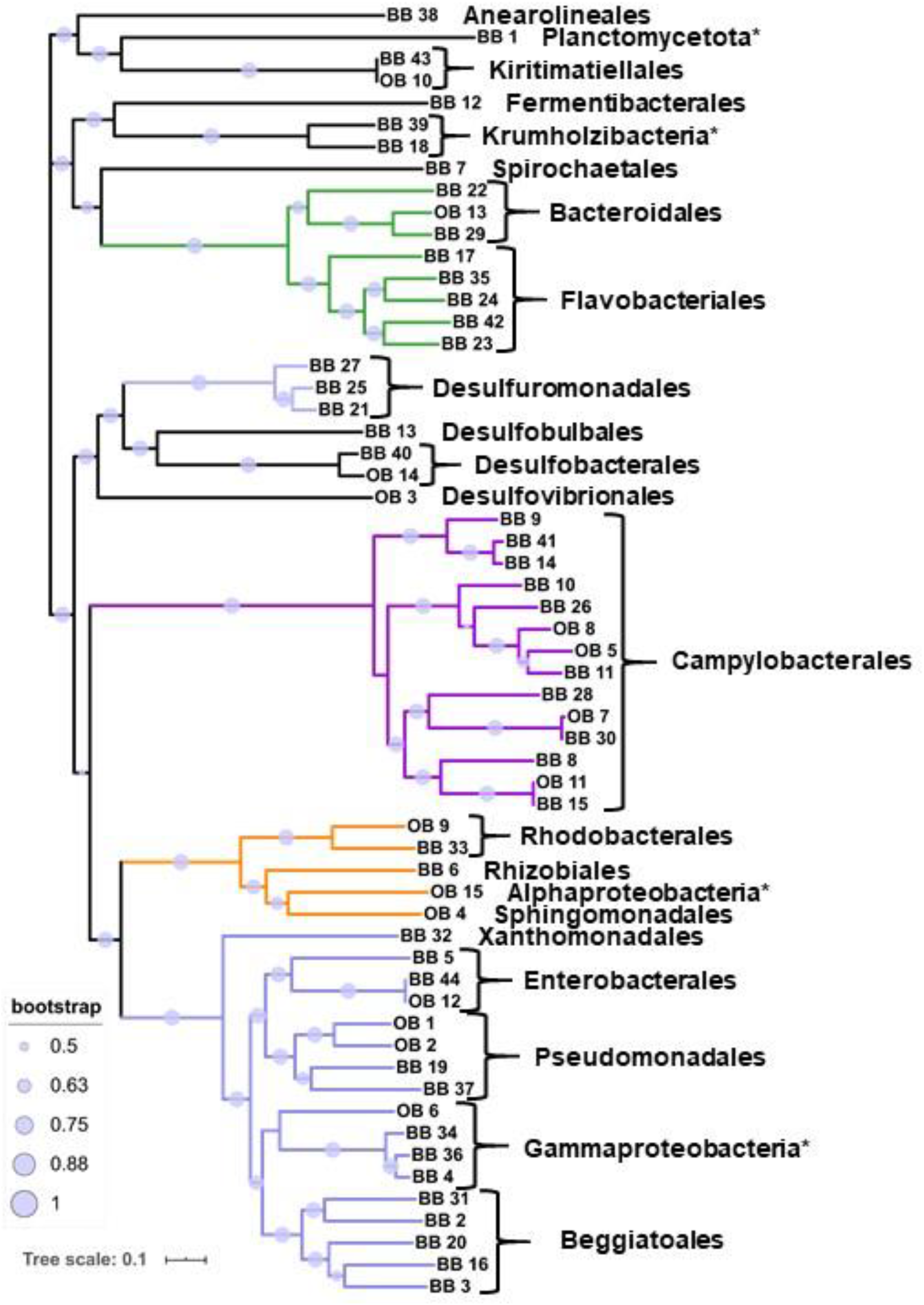
Phylogenomic maximum likelihood tree of the obtained 59 high-quality MAGs from *Osedax*-associated bone microbiome (OB) and bone surface-associated biofilm (BB). Bootstrap values greater than 0.5 are displayed as circles on the branches. The five most common bacterial classes are colored (green – Bacteroidia, light purple – Desulfuromonadia, purple – Campylobacteria, orange – Alphaproteobacteria and blue – Gammaproteobacteria). Order level identifications are listed and MAGs for which only class level identification could be inferred are marked with an asterisk.

Besides a large number of SOB, eight MAGs related to SRB were identified that are affiliated to the families Desulfobulbaceae (also SOB), Desulfobacteraceae, Geopsychrobacteraceae and Desulfovibrionacaeae. Moreover, they are prevalently associated with the free-living community attached to the bone surface in this study. Sulfate, tetrathionate or thiosulfate can serve as electron acceptors and/or donors and gene markers for all pathways are present in the metagenomes (Figure 2). Microbial sulfidogenesis on the bone surface or the surrounding sediments can feed the thiotrophic community and therefore accelerate the demineralization process. The generated sulfide is known to quickly react with iron, blackening the bone surfaces with insoluble iron sulfide (77). In our incubation experiments, blackening occurs preferentially on the epiphysis, which is also where complex white/pink microbial mats are forming over time (Supplementary figure S1B). However, from our analysis SRB seem unable to degrade large complex molecules. This is supported by the lack of bone-degrading enzymes herein investigated, such as: S8/S53 peptidases, mannosidases, sialidases, fucosidases and α-galactosidases. SRB are likely dependent on the generation of simple organic compounds produced as metabolites by fermenters or aerobic organotrophic bacteria of the wider bone microbiome. The bone dissolution driven by sulfur geomicrobiology relies on other specialized members of the community to degrade the organic matrix and to fuel the acid generation.

**Figure 2:**
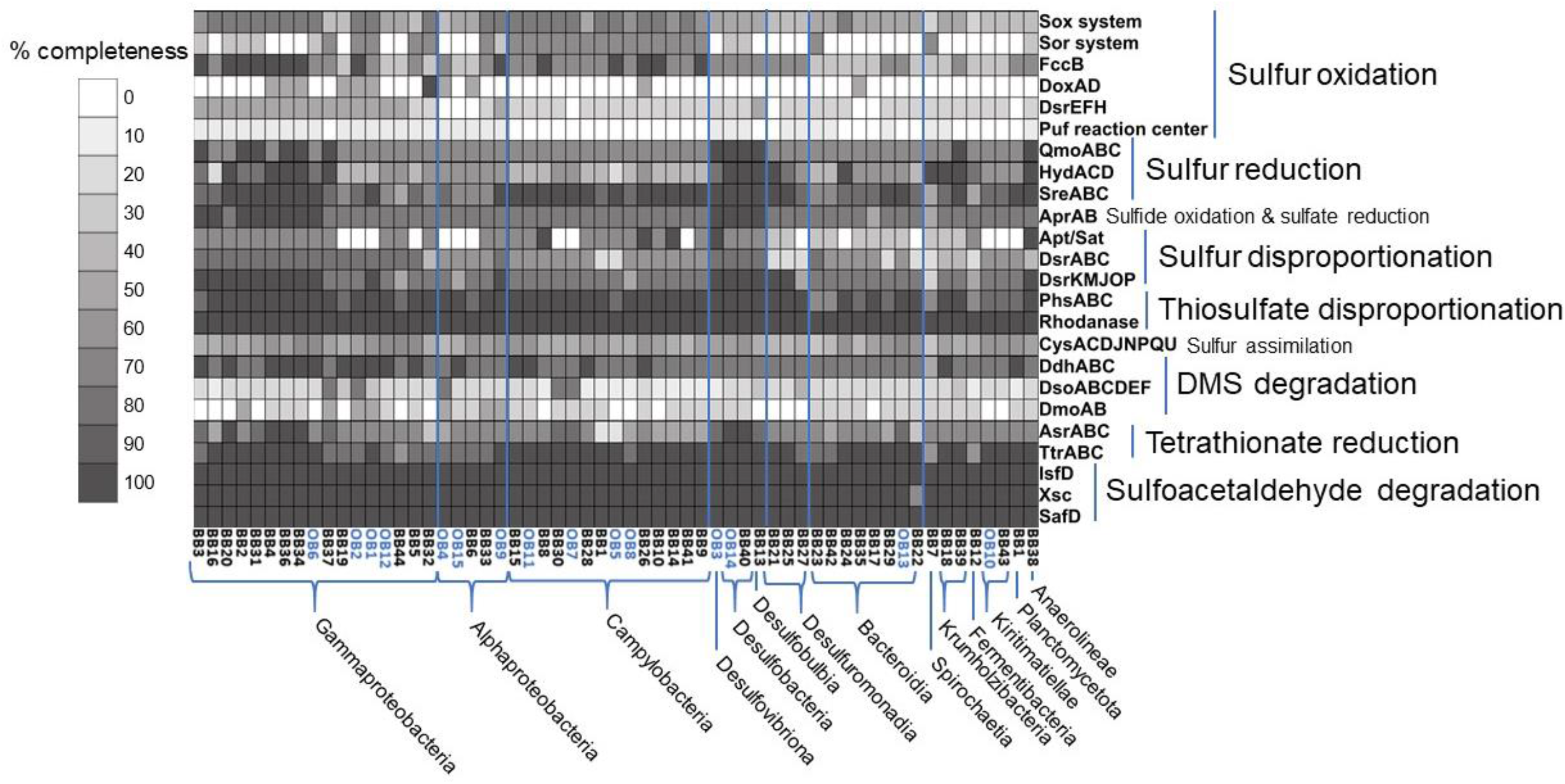
Whole genome metabolic pathway comparison for genes of the sulfur metabolism. Analysis was done with MEBS (37) and MAGs were phylogenetically grouped according to GTDBTk pipeline (36). The grey scale represents the completeness of a given pathway or multienzyme system shown in the heatmap for each MAG. The OB MAGs are highlighted in blue. The “Sox system” constitutes of the *sox*XYZABCD genes and the “Sor system” of *sor*ABDE.

#### Degradation of organic compounds via peptidases, glucosidases and oxidases

Once the inorganic hydroxyapatite is removed, an array of different enzymes is required to digest the various organic bone components. Bacteroidia appear to be especially remarkable in this respect and represent the third most abundant taxon. Eight high quality MAGs could be reconstructed, seven of them from the bone surface metagenome. Bacteroidia, and especially the family of Flavobacteriaceae, are known to be versatile degraders of polysaccharides like agar (78), chitin (79), ulvan (80), alginate (81), carrageen (82), cellulose and xylanose (83) and polypeptides like elastin (84), spongin (85) and others. The recently described Marinifilaceae family (86) includes isolates that are reported to present xylanase activity (87). Despite the discrepancy between abundance versus reconstructed genomes, the Bacteroidia MAGs appear to be the most versatile order of the investigated MAGs in respect to their richness in bone degrading enzymes (Figure 4), and all were predicted to possess the gelatin hydrolysis trait (Figure 3). They were also the only MAGs containing sialidases (COG4409) and α-galactosidases (Pfam16499) (Figure 4). Due to this taxa-specific trait and their presence limited to the bone-surface associated microbiome, we hypothesize that Bacteroidia play a pivotal and specialized role in the free-living community via the degradation of specific organic bone components.

**Figure 3:**
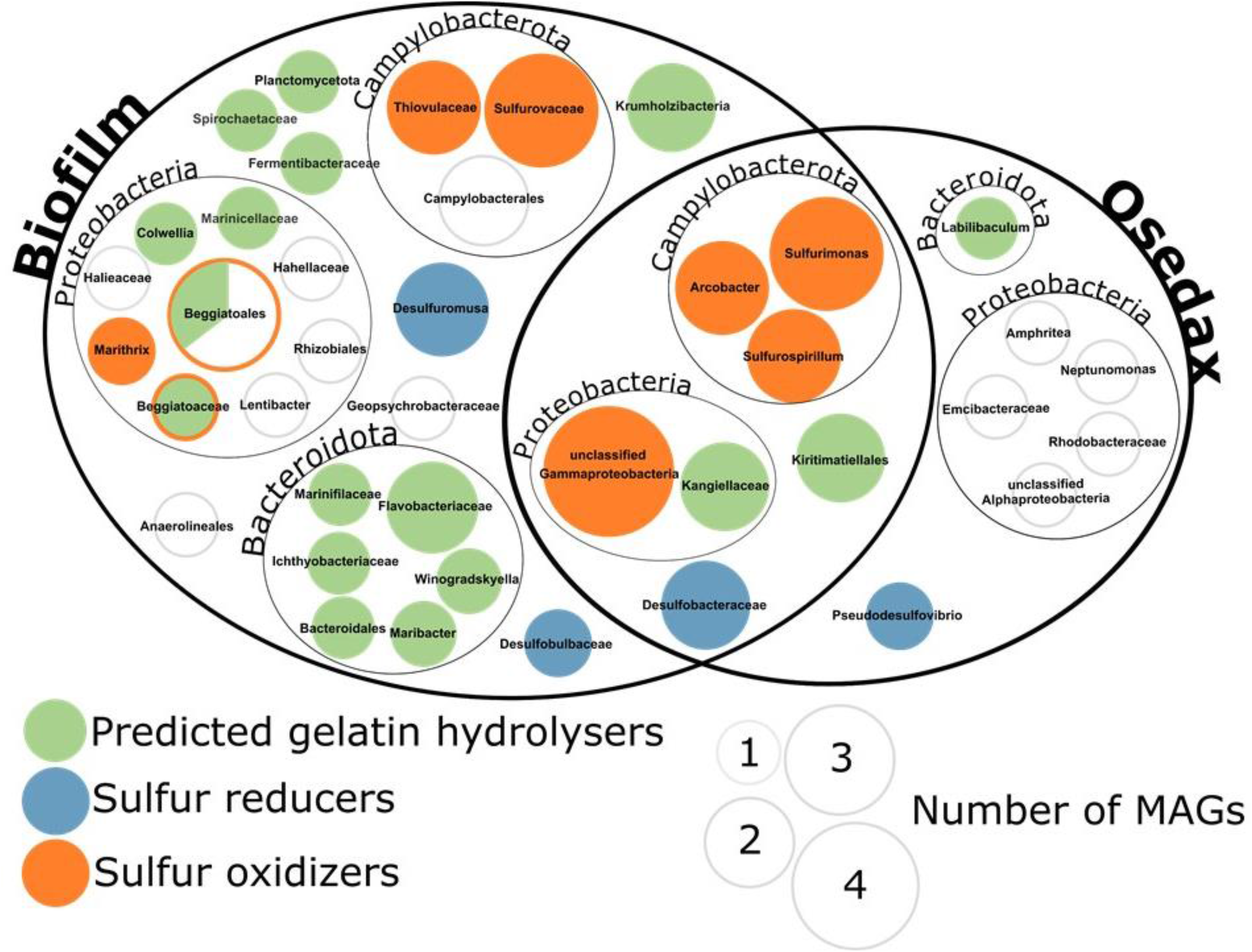
Abundance and taxonomic affiliation of predicted gelatin hydrolysers (green) (Traitar) in *Osedax* and biofilm derived MAGs. Additionally, sulfur reducers (blue) and sulfur oxidizers (orange), (MEBS) are shown, while white indicates absence of these traits. MAGs are displayed at the deepest taxonomic classification obtained. The size of the circles reflects the number of MAGs within each clade.

**Figure 4:**
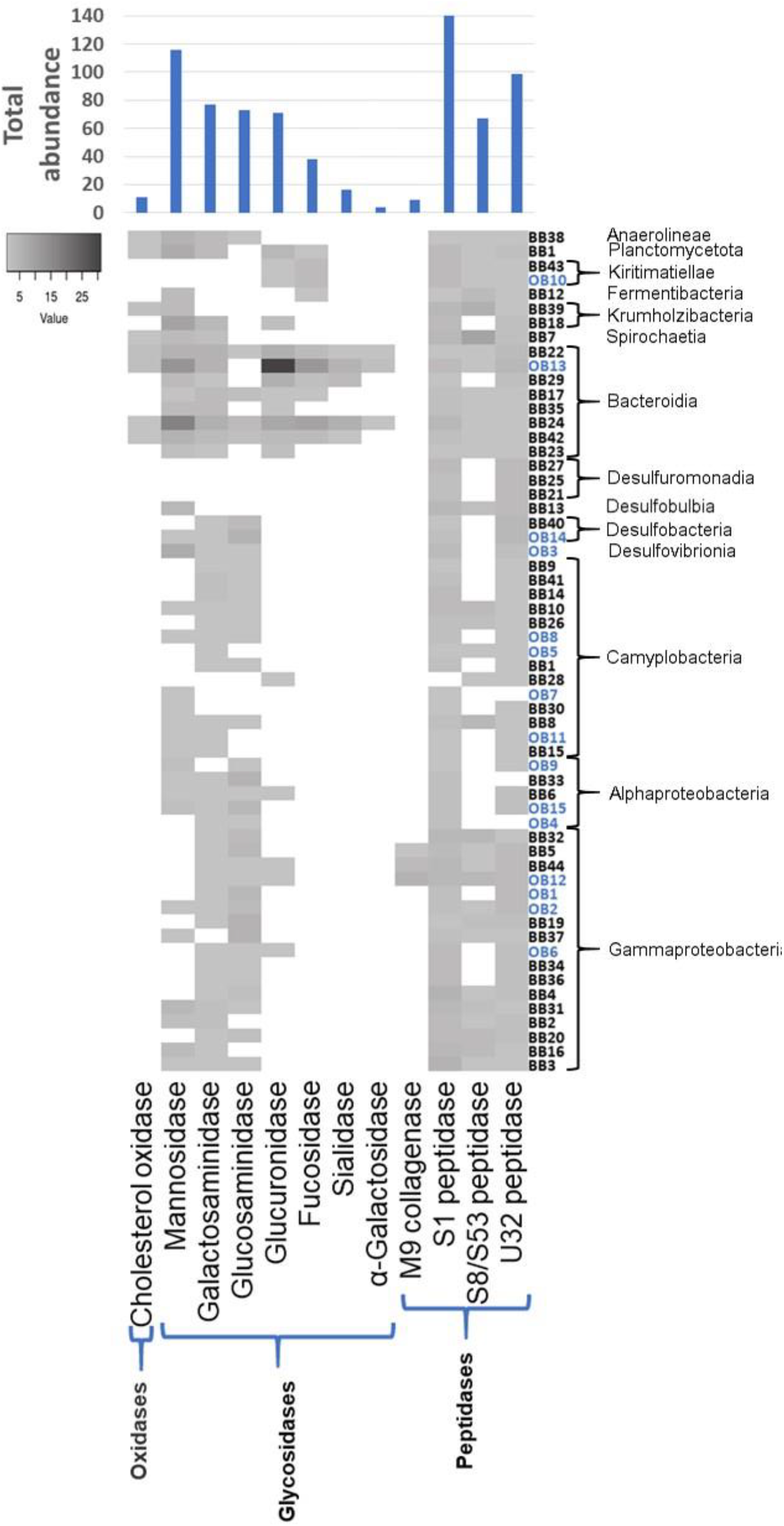
Abundance heatmap of the 12 investigated enzyme COG classes in the 59 bone degradome MAGs. The MAGs are arranged according to their taxonomic affiliation. The absolute abundance of each enzyme COG class is depicted in the diagram on top of the heatmap. The OB MAGs are highlighted in blue.

Differential microbial colonization of the spongy cancellous bone tissue over the cortical compact bone has also been observed in the terrestrial environment and has been related to easier access to the red marrow (88) although a priming effect linked to the differential composition of the bone cannot be ruled out. Complex microbial mats form preferentially on the epiphysis of the long bones and this area is normally covered with hyaline cartilage (89) which was not removed before deployment. Cartilage is a related connective tissue made of nonfibrous type II collagen and a sulfated-proteoglycan matrix rich in N-acetyl-galactosamine and glucuronic acid residues. This would explain the abundance of alpha-galactosidases, N-acetyl glucosaminidase and glucuronidases. Moreover, other groups such as Kiritimatiellales (PVC superphylum) are known marine anaerobic saccharolytic microbes specialized in degrading sulfated polymers that we find in this environment (90).

#### Collagen degradation by Gammaproteobacteria

Peptidases and especially M9 collagenases are of special interest for the degradation of the proteinogenic compounds within bone, as they are able to degrade collagen, the main source of carbon in this environment. The class γ-proteobacteria is comparatively enriched in these enzymes and it is the best represented class in the dataset, with 17 MAGs. Of particular interest are the MAGs affiliated to the order Enterobacterales (two MAGs of the families Kangiellaceae and one Alteromondaceae). They possess the gelatin hydrolysis trait (Figure 3, MAGs BB5, BB44 and OB12), have a high number of S1 and U32 peptidases, and are the only MAGs with M9 collagenases. The *Colwellia* MAG BB5 is particularly remarkable as it contains an entire gene cluster dedicated to collagen utilization (Figure 5A). The collagen degradation gene cluster comprises at least 15 different genes, including a M9 collagenase, a PepQ proline dipeptidase, an aminopeptidase YpdF, several transporters, epimerase, isomerases and others. The gene cluster encodes nearly the entire pathway necessary to unwind and hydrolyze triple-helical collagen, transport and uptake of collagen oligopeptides into the cell and utilization of its main components, mainly hydroxyproline and proline, for energy production via the TCA cycle and/or the urea cycle or for polyamine biosynthesis (Figure 5B). Accessory genes for an alternative catabolic route of proline to glutamate are also encoded elsewhere in the genome (P5CDH). This kind of functional condensation for collagen utilization has not been described before in *Colwellia* or elsewhere. Interestingly, *Colwellia* bacteria are also one partner in a dual extracellular symbiosis with sulfur-oxidizing bacteria in the mussel *Terua* sp. ‘Guadelope’, retrieved from a whale fall in the Antilles arc and supposedly involved in the utilization and uptake of bone components (91). A cluster of functionally related genes was found in the publicly available genomes of *Col·wellia piezophila* and *Colwellia psychrerythraea.* However, the gene cluster described for MAG BB5 contains several supplementary features, like a rotamase, a 1-pyrroline-4-hydroxy-2-carboxylate deaminase and a Zn/Fe binding domain protein potentially attributed to collagen utilization, which are absent in the published genomes (Figure 6). Moreover, the gene cluster contains regulatory elements like the PutR regulator and stringent starvation proteins known to be activated under acid stress or amino acid starvation conditions in *Escherichia coli* (92). This supports our hypothesis that other members of the microbial community need to dissolve the bone calcium phosphate via acid secretion, before collagen and other organic bone compounds can be accessed. In summary the publicly available gene clusters lack regulatory elements to switch on the collagen utilization pathway under ‘bone degrading’/acidified conditions and misses key enzymes to exploit collagens key components proline (rotamase missing to transverse D-proline to L-proline) and hydroxyproline (pyrroline-4-hydroxy-2-carboxylate deaminase missing that breaks down 1-pyrroline-4-hydroxy-2-carboxylate to alpha-ketoglutarate semialdehyde).

**Figure 5:**
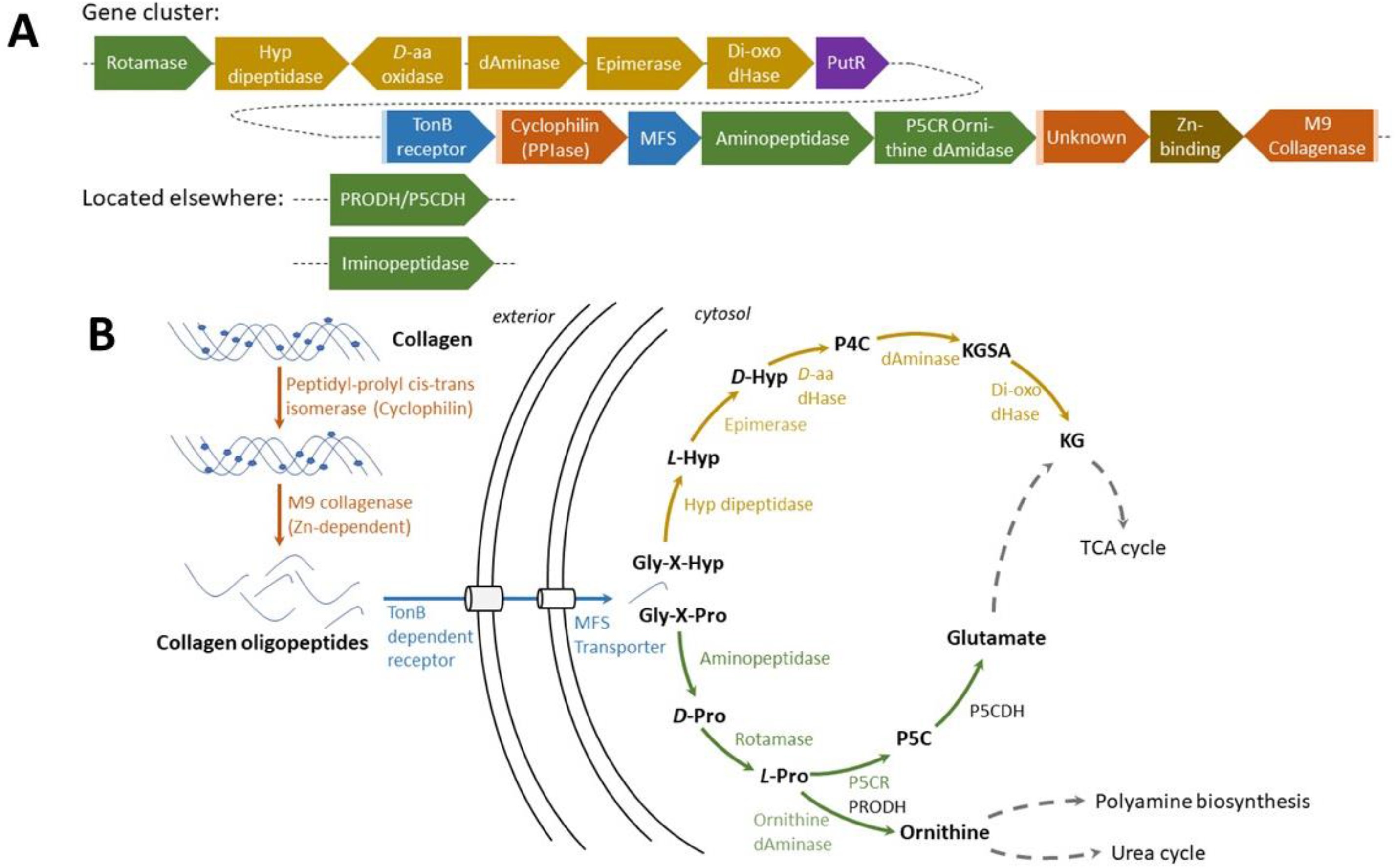
Collagen utilization pathway scheme in MAG BB5. A) The gene cluster in BB5 spans approximately 21 kb, comprising 15 genes for collagen utilization, each color-coded respective to its functional group: orange for collagen hydrolysis, blue for uptake and transport, green for proline (Pro) utilization, ocher for hydroxyproline (Hyp) utilization and brown for unknown function. The purple box is indicative of a signal-peptide for secretion. B) Metabolic model for collagen utilization in *Colwellia* BB5. Arrows and genes are color-coded in the same functional groups as in A. Dashed arrows point to a major metabolic pathway. Metabolite abbreviations: P4C (1-pyrroline 4-hydroxy-2-carboxylate), KGSA (alpha-ketoglutarate semialdehyde), KG (alpha-ketoglutarate), P5C (1-pyrroline-5-carboxylate). Enzyme abbreviations: D-aa dHase (D-hydroxyproline dehydrogenase), dAminase (pyrroline-4-hydroxy-2-carboxylate deaminase), dioxo dHase (KGSA dehydrogenase), P5CR/ornithine dAminase (bifunctional 1-pyrroline-5-carboxylate reductase/ornithine cyclodeaminase), PRODH (proline dehydrogenase), P5CDH (pyrroline-5-carboxylate dehydrogenase).

**Figure 6:**
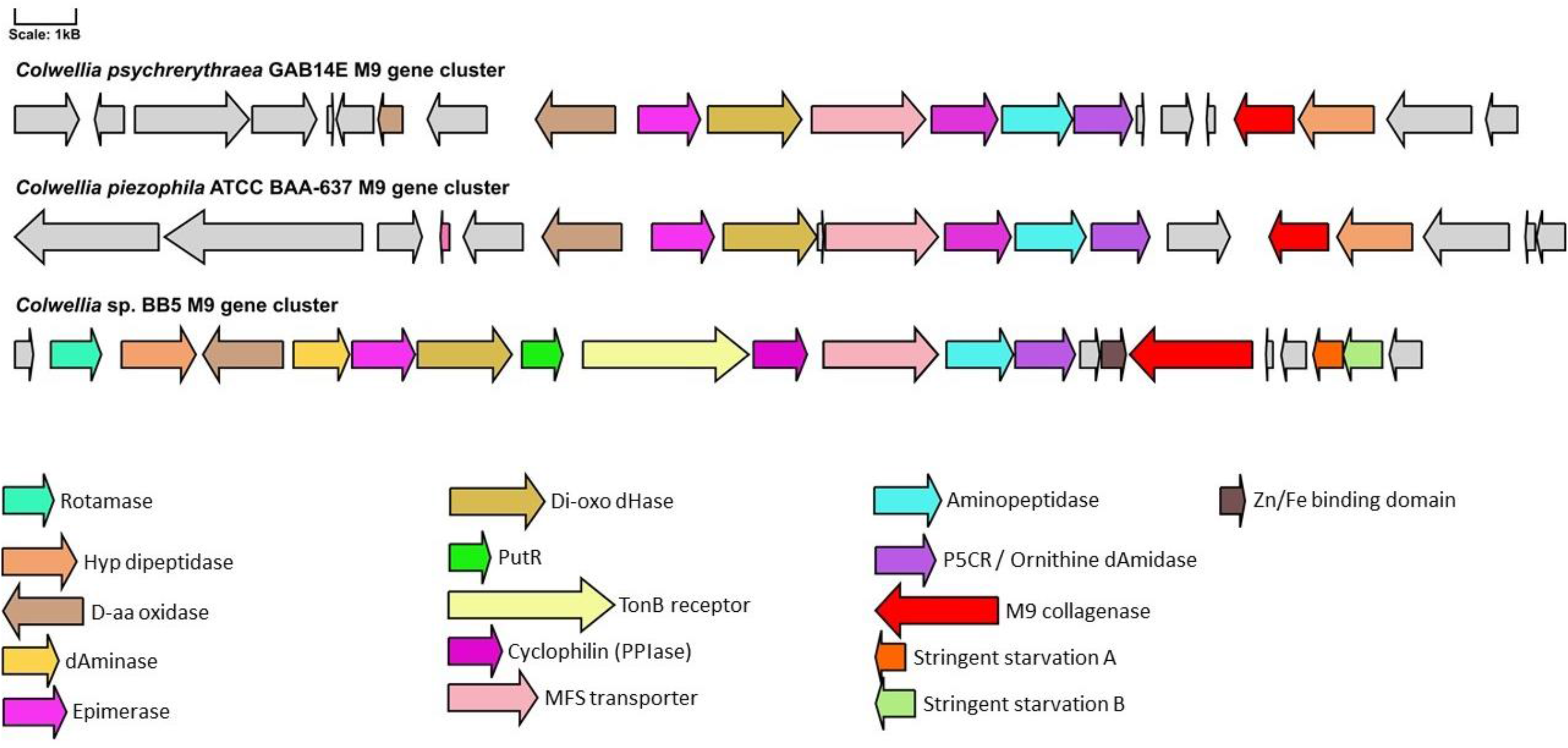
Conservation between M9 collagen degradation gene clusters in *Colwellia psychrerythraea* GAB14E, *Colwellia piezophila* ATCC BAA-637 and the MAG *Colwellia* BB5 drawn at scale. dHase, dehydrogenase; PPIase, peptidyl-prolyl cis trans isomerase; Hyp, D-aa, dAminase etc. Color coding and gene names are indicated.

#### Bone degradation – a complex microbial community effort

The marine bone microbiome is a complex assemblage of various bacterial classes that requires the synergistic action of many different interwoven enzymatic reactions to access the recalcitrant bone material for its nutritional resources. A scenario how we envision the orchestration of this complex process is depicted in Figure 7. The primary requirement in utilizing organic bone compounds is likely the dissolution of mineralized calcium phosphate (hydroxyapatite) by acidification, which can potentially be performed via proton release by a versatile community of sulfur-oxidizing (SOB) γ-proteobacteria (mainly *Beggiatoa*-like), Campylobacterales *(Sulfurimonas, Sufurospirillum, Sulfurovum),* Desulfobubales and α-proteobacteria (Figure 7-I). This acidification via thiotrophy may be fueled by sulfur-reducing bacteria (SRB), like Desulfobacteraceae, Geopsychrobacteraceae, *Pseudodesulfovibrio,* creating a sulfur biogeochemical loop between SRB and SOB (Figure 7-II). Once the organic compounds (collagen, fatty acids, proteins, peptidoglycans) are accessible, the Bacteroidia (Flavobacteriaceae and Marinifiliaceae) and γ-proteobacteria (Alteromonadaceae and Kangiellaceae) may become the main protagonists (Figure 7-III and IV). These Bacteroidia are especially rich in bone degrading enzymes, but importantly the γ-proteobacteria are the only members identified with M9 collagenases and *Colwellia* contains an entire gene cluster dedicated to collagen degradation (Figure 5). Herein we disentangled the potential functional roles of specialized members of the bone-degrading microbial community, which together make bone-derived nutrients accessible – not only to themselves, but also to generalists within the bone microbiome. We posit that Flavobacteriales and Enterobacterales are the most promising candidates for novel enzyme discovery, as they display the most versatile sets of bone degrading enzymes.

**Figure 7:**
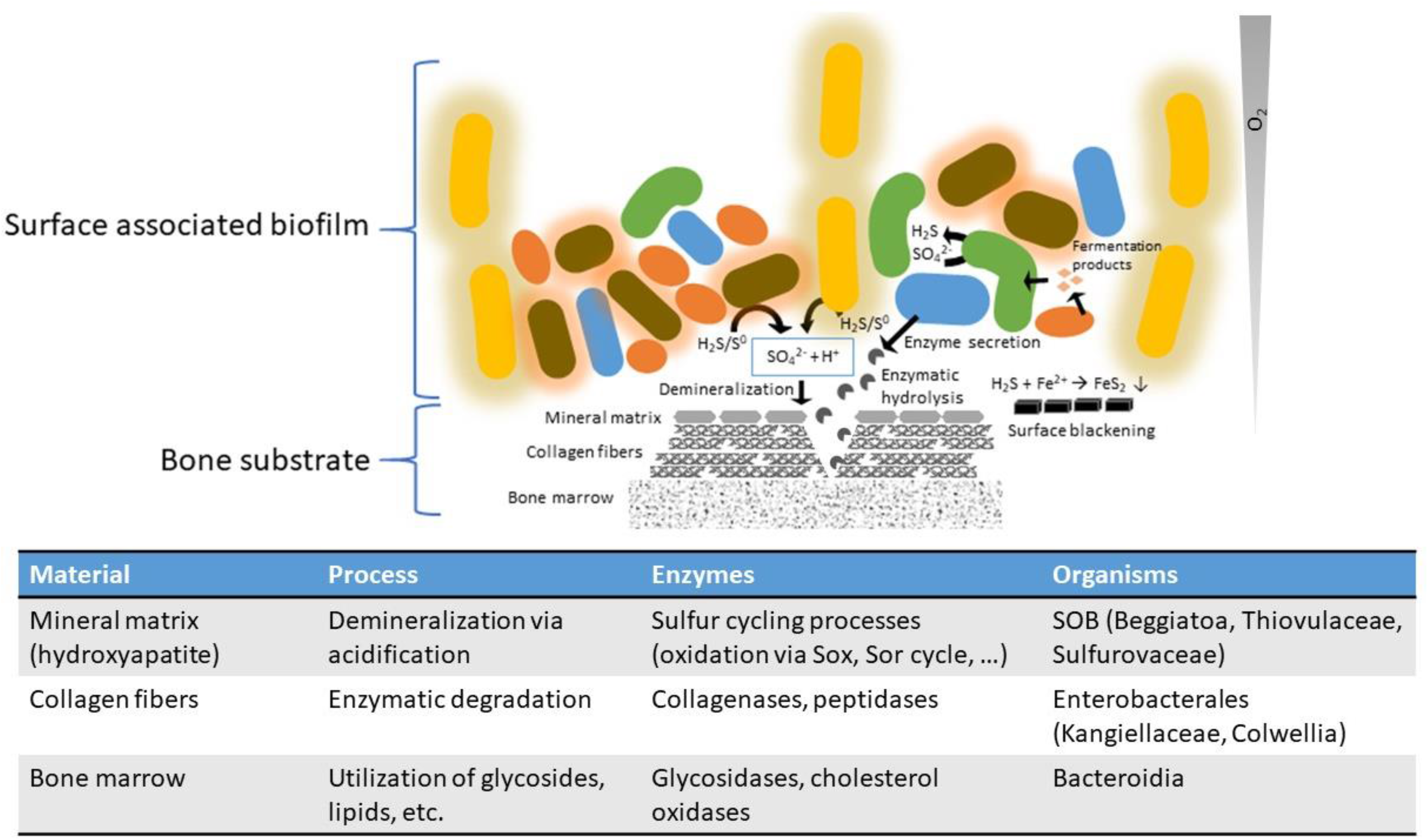
Hypothesis of the interplay in the marine bone microbiome and degradome. Sulfuroxidizing bacteria (SOB, shown with a halo) convert elemental sulfur and H_2_S into sulfate and protons that lead to an acidification and therefore bone demineralization. Sulfate reducing (SRB, green) and sulfur disproportioning bacteria produce H_2_S from sulfate. Enterobacterales and other especially Gammaproteobatceria secret collagenases to degrade collagen. Bacteroidia and other bacteria secret gylcosidases and other enzymes to hydrolyze the organic bone components (glycosides, esters, lipids). This exemplifies a bone demineralization loop that fuels itself as long as sulfur is available and degrades the organic bone components in the process.

## Materials and Methods

### Sample collection

Bone material after manual meat deboning was kindly provided by a local slaughterhouse operated by Norilia (Norway). Since deboning does not completely eliminate the animal tissue attached to the bone, some remains were still present. Therefore, in order to avoid bacterial colonization and decomposition, all bone material was kept at −20 C until deployment. Four sets of turkey thigh bones and one bovine lower leg bone were placed in a crab trap and deposited at the bottom of Byfjorden (60,397093N; 5,301293E) close to Bergen, Norway, at a depth of 69 m and approximately 150 m off shore in May 2016, incubated for nine months and retrieved using a small ROV (Table 1). The material was transported to the lab in styrofoam boxes for either processing within two hours (bone surfaces) or for prolonged incubation in seawater aquaria and subsequent dissection of *Osedax* worms. The meatless bone surfaces were scraped with a sterile scalpel for microorganisms and *Osedax mucofloris* specimens were extracted from the bone using sterile scissors and forceps. Their root tissue was dissected from the body, rinsed in sterile 70% (v/v) sea water and preserved in storage solution (700 g/l ammonium sulfate, 20 mM sodium citrate and 25 mM EDTA, pH 5.2) and stored at −70 °C until further processing.

### DNA extraction and sequencing

DNA was extracted from 10 to 50 mg sample of either scrapped biofilm or *Osedax* root tissue, using the Qiagen AllPrep DNA/RNA Mini Kit according to the manufacturer’s instructions with cell lysis by a bead beating step in Lysing Matrix E tubes (MP Biomedicals) in a FastPrep homogenizer (MP Biomedicals) with a single cycle of 30s at a speed of 5500 rpm. The obtained DNA was quantified and quality controlled using a NanoDrop2000 (ThermoFisher Scientific) and a Qubit fluorometer 3.0 (ThermoFisher Scientific). The obtained DNA concentrations ranged from 11,9 ng/μl to 166 ng/μl according to Qubit readings. The DNA libraries were prepared using the Nextera DNA library Prep Kit according to the manufacturer’s guidelines. 50 ng DNA was used for the preparations. In brief, the DNA was fragmented by the Nextera transposome at 55°C for 5 min and barcoded adapters were added in a 5 cycle PCR amplification. The resulting libraries, including ~140 bp of adapter had an average fragment size of 436 bp (± 112 bp) and were sequenced on an Illumina HiSeq4000 platform (150-bp paired-end reads) at the Institute of Clinical Molecular Biology (IKMB), Kiel University, Germany.

### Metagenomic read profiling

Illumina raw reads were quality trimmed and adapters were removed with Trimmomatic version 0.36 (93). The quality filtered reads were used individually and combined with respect to their sample source (either *Osedax-associated* or bone surface biofilms) to profile the taxonomic origin of the reads with Kaiju (35).

### Metagenomic assembly, binning, taxonomic identification, ORF prediction and annotation

For each sample type *(Osedax mucofloris* and bone surface biofilm communities), the quality filtered metagenomic reads (Trimmomatic version 0.36) were co-assembled with SPAdes v3.12 (94) for kmers 21, 33, 55, 77 and 99, with the metaSPAdes-assembler option enabled (see Supplementary table S1 for read counts). Binning was conducted on the resulting assemblies using the MetaWRAP pipeline (version 1.0.1) (95). This pipeline combines three different implemented binning methods, CONCOCT (96), MaxBin2.0 (97) and metaBAT2 (98), to retrieve high quality MAGs. We only considered high-quality MAGs with >90% completeness and <10% redundancy for further analyses. CheckM was used for quality assessment of the assembled genomes (99), and GTDB-Tk version 0.1.3 (36) was used for taxonomic identification, coupled with an estimate of relative evolutionary divergence (RED) to their next common ancestor. RED is a normalization method to assign taxonomic ranks according to lineage-specific rates of evolution, based on branch lengths and internal nodes in relation to the last common ancestor calculated by GTDBTk. Open reading frames (ORF) of the obtained MAGs were predicted with Prodigal version 2.6.3 (100). Predicted ORFs were annotated using eggNOG-mapper v1 (101) with eggNOG orthology data version 4.5 (102). Additionally, the MAGs were annotated and metabolic models were calculated using the RAST (rapid annotation using subsystem technology) server (103, 104). The MAGs were further investigated for the presence or absence of major metabolic pathways and phenotypic microbial traits based on their genomic sequences using MEBS (multigenomic entropy based score, version 1.2) (37) and Traitar (40). MEBS is a software package used here to detect genes related to sulfur metabolism, this was done by providing protein fasta files of the high-quality MAGs, that are annotated by MEBS with Interproscan (105) and are then searched with HMM profiles for genes related to sulfur metabolism. The sulfur metabolism related genes investigated by MEBS are based primarily on the MetaCyc database (106). Traitar is a software package that can predict 67 phenotypic traits from a genome sequence. In brief, the analysis is based on known phenotypic traits of 234 bacterial species and infers from their genome Pfam families that are either present or absent in a specific trait. In this manuscript the traits gelatin hydrolysis and H_2_S production are of interest and these are based the presence of 70 and 43 and absence of 51 and 22 Pfam families respectively. Phylogenomic trees were calculated with FastTree (107) as maximum likelihood trees and visualized with iTOL (108, 109) and heatmaps were visualized with Heatmapper (110). Gene cluster maps were drawn with Gene Graphics (111). Signal peptides were predicted with SignalP-5.0 server using nucleotide sequences to predict the presence of Sec/SPI, Tat/SPI and Sec/SPII signal peptides in a given sequence (112).

### Enzyme profiling

Based on the organic composition of bone matrix, we hypothesized twelve enzyme families to be necessary for its degradation. Accordingly, the following enzymes were selected for in-depth studies: (i) M9 collagenases (pfam01752), S1 peptidases (COG0265), S8/S53 peptidases (pfam00082) and U32 proteases (COG0826) which hydrolyze peptide bonds in collagen and glycoproteins; (ii) sialidases (COG4409), β-D-glucuronidases (COG3250), β-N-acetyl-D-glucosaminidases (COG1472), α-N-acetylgalactosaminidases (COG0673), α-galactosidases (pfam16499), fucosidases (COG3669) and mannosidases (COG0383) which cleave glycosidic lingakes, and (iii) cholesterol oxidases (COG2303) which degrade lipids such as cholesterol. One reference database for each of these families was generated using the NCBI repository, based on sequences from 287 M9 collagenases, 4453 S1 peptidases, 3237 S8/S53 peptidases, 3653 U32 proteases, 267 COG4409, 873 COG3250, 1274 COG1472, 6140 COG0673, 279 COG3669, 206 COG0383 and 1119 COG2303. The databases included the closest protein homologs of all protein families of interest for bone-degradation, and at least one representative sequence from all taxonomic groups (containing such enzymes) was represented. The reference databases were used to generate Hidden Markov Model (HMM) profiles for each enzyme family with HMMER version 3.1b1 (113) using the *hmmbuild* option after an alignment of each sequence set was built with Clustal W version 2.1 (114). The MAGs were screened for the twelve enzyme families of interest using the generated HMM profiles using HMMer version 3.1b1 with the *hmmsearch* option and a bitscore threshold of 100.

### Tara Oceans comparison

This comparison is based on the work of Meren *et al.* (2018), who binned 93 metagenomes generated from the Tara Oceans project (115). The metagenomes represent 61 surface water samples and 32 samples from the deep chlorophyll maximum layer of the water column. They generated 957 non-redundant high-quality bacterial, archaeal and eukaryotic genomes. The 957 MAGs were here reanalyzed with GTDB-Tk and the generated HMM profiles as previously described. 832 MAGs were identified as of bacterial origin and included in the comparison. The Tara Oceans MAGs were generated with a quality threshold of >70% completion, therefore the bone metagenomes were rebinned (>70% completion, <10% redundancy) for better comparison to avoid bias due to the higher threshold used for functional analysis in other parts of this manuscript.

## Data availability

The raw sequencing reads have been deposited in the sequence read archive (SRA) of NCBI under the BioProject ID PRJNA606180 and with the BioSample accession numbers SAMN14086998 (A5), SAMN14087000 (A9), SAMN14087001 (A9n), SAMN14087003 (B4), SAMN14087005 (D1), SAMN14087006 (D2), SAMN14087007 (I1) and SAMN14087008 (I3).

The 59 high-quality MAGs analyzed in this study were deposited in the NCBI database as well, as part of the BioProject ID PRJNA606180, the BioSample accession numbers are SAMN16086327-SAMN16086385 (Biofilm MAGs 1-44 SAMN16086327-SAMN16086370 and *Osedax* MAGs 1-15 SAMN16086371-SAMN16086385).

## Authors contributions

T. D., E.B., A.G.M. and G.E.K.B. developed the conceptual idea, conducted the sampling of the source material, contributed to content, reviewed and edited the manuscript. S.S.C., E.B. and M.F. identified enzymes of interest, constructed the reference databases and aided in result interpretation. E.B., B.S., S.F. and A.G.M. conducted bioinformatic analyses and E.B., B.S. and U. H. wrote the manuscript.

## Acknowledgements

To the memory of Prof. Hans Tore Rapp and his effort on characterizing *Osedax mucofloris.* We acknowledge financial support of ERA-NET Marine Biotechnology (GA no.: 604814) funded under the FP7 ERA-NET scheme and nationally managed from the German Federal Ministry of Education and Research and Norwegian Research Council. M.F. acknowledges the support of the grants PCIN-2017-078 (within the Marine Biotechnology ERA-NET) and BIO2017-85522-R from the Ministerio de Economía y Competitividad, Ministerio de Ciencia, Innovación y Universidades (MCIU), Agencia Estatal de Investigación (AEI), Fondo Europeo de Desarrollo Regional (FEDER) and Competiveness. Additional funding was received from the Norwegian Biodiversity Information Centre (PA 809116 knr. 47-14). We thank Hans T. Kleivdal for early developments of the concept. We thank Norilia AS for supplying bone residue material for the field work and ROV AS for providing underwater services during the deployment and sampling campaign. The authors acknowledge Kira S. Makarova at the National Center for Biotechnology Information (NCBI, Bethesda MD, USA) for help during the design of the reference database of enzyme families relevant to bone degradation.

## Supplementary material

### Supplementary tables

**Supplementary table S1:**
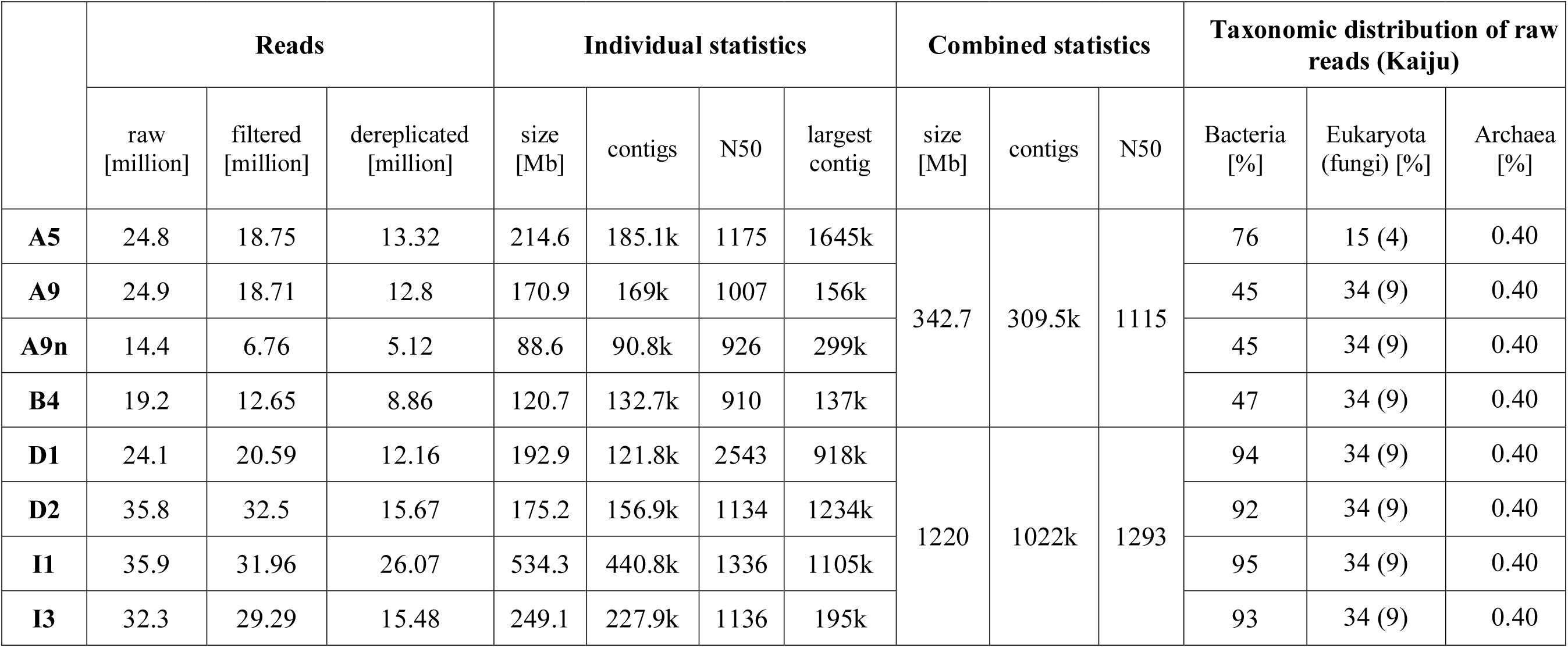
Metagenome statistics

**Supplementary table S2:**
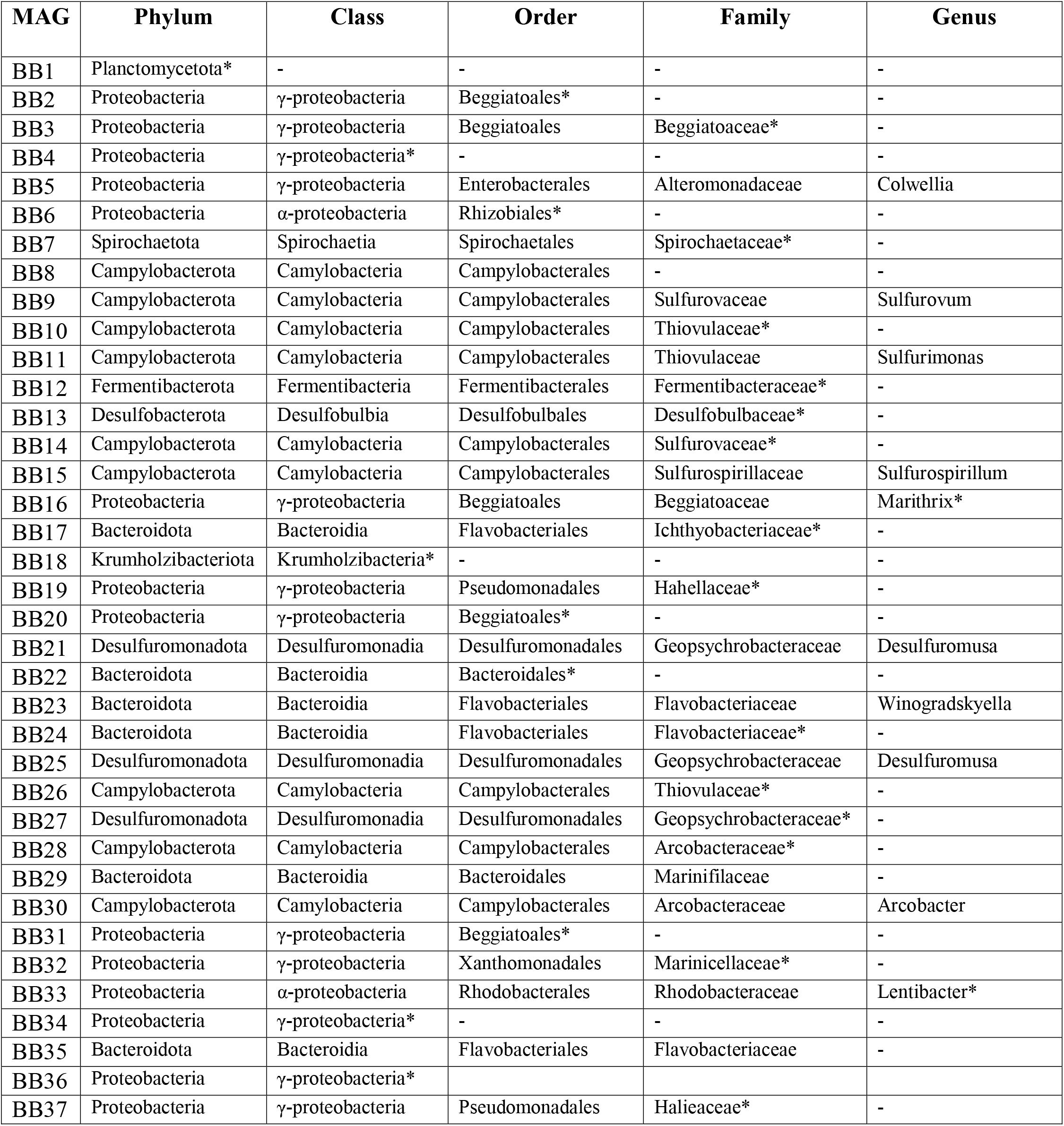
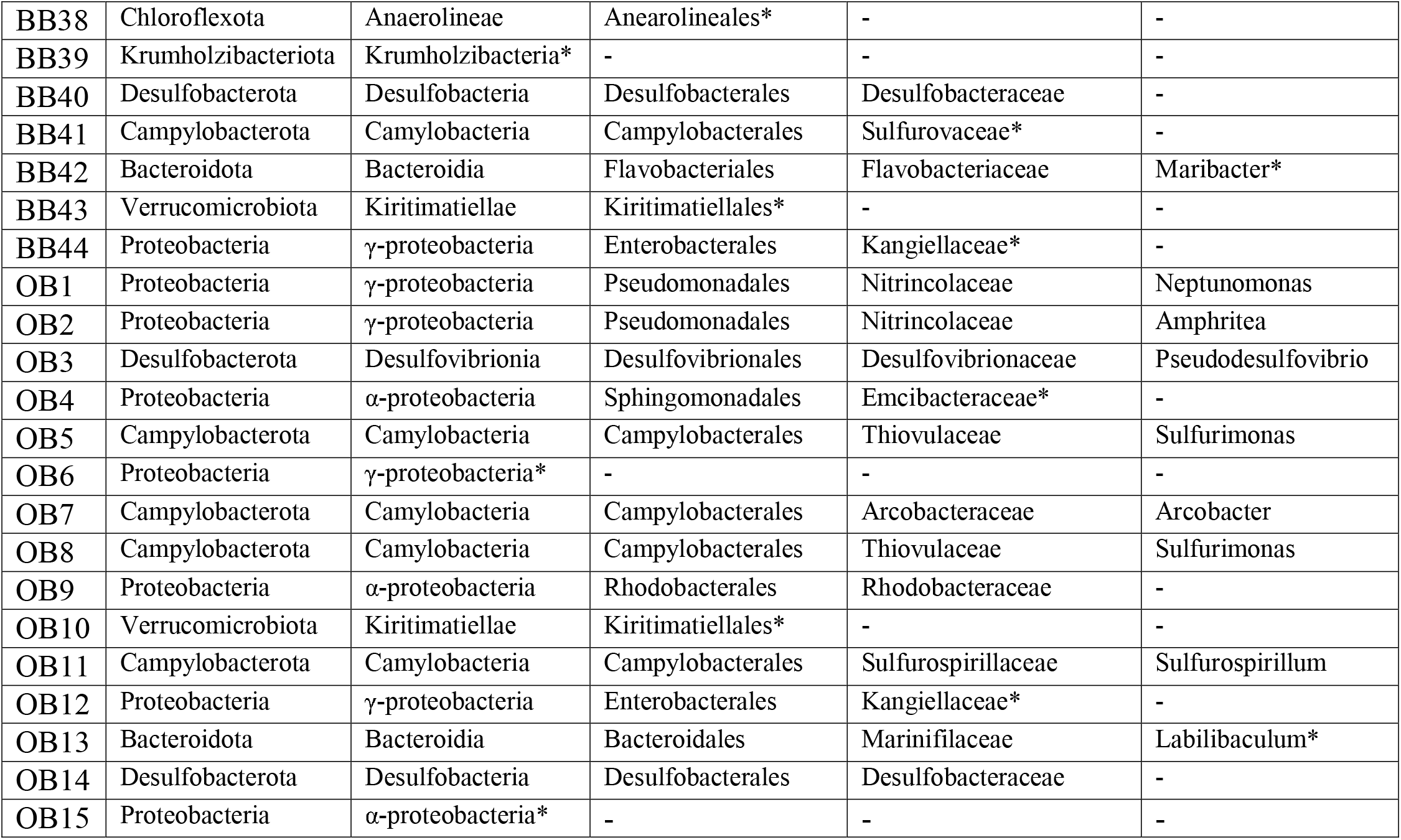
Taxonomic affiliation of MAGs according to Parks *et al.* (2018) and taxonomic novelty identified by RED (*) (36).

**Supplementary table S3:** MAG sequence data. All MAGs labeled ‘OB’ were retrieved from Osedax samples and all MAGs labeled with ‘BB’ from bone surface biofilms.

| MAG | Genome size [Mbp] | Longest contig [kbp] | N50 [kbp] | No. of contigs | Predicted genes | GC content [%] | No. of bone-degrading Enzymes |
| --- | --- | --- | --- | --- | --- | --- | --- |
| BB1 | 4.68 | 155 | 429 | 38 | 3723 | 44.50 | 24 |
| BB2 | 4.18 | 853 | 248 | 108 | 3538 | 44.10 | 13 |
| BB3 | 4.65 | 75 | 23 | 362 | 4126 | 37.70 | 11 |
| BB4 | 5.18 | 1.050 | 198 | 68 | 4705 | 38.80 | 13 |
| BB5 | 3.26 | 82 | 13 | 395 | 3018 | 36.10 | 16 |
| BB6 | 2.77 | 72 | 19 | 244 | 2783 | 40.10 | 10 |
| BB7 | 4.87 | 78 | 15 | 489 | 4636 | 40.10 | 22 |
| BB8 | 3.08 | 89 | 18 | 413 | 3332 | 32.90 | 11 |
| BB9 | 2.16 | 126 | 36 | 159 | 2259 | 36.80 | 5 |
| BB10 | 3.01 | 137 | 35 | 142 | 2950 | 36.20 | 11 |
| BB11 | 2.39 | 21 | 3 | 809 | 2958 | 35.40 | 6 |
| BB12 | 3.24 | 329 | 141 | 127 | 2929 | 45.10 | 10 |
| BB13 | 4.24 | 16 | 3 | 1688 | 4778 | 48 | 16 |
| BB14 | 2.32 | 74 | 33 | 103 | 2265 | 34.20 | 8 |
| BB15 | 2.68 | 415 | 191 | 33 | 2695 | 30.70 | 5 |
| BB16 | 3.09 | 481 | 237 | 22 | 2867 | 34.10 | 13 |
| BB17 | 3.64 | 81 | 16 | 327 | 3171 | 32.30 | 13 |
| BB18 | 4.43 | 267 | 114 | 176 | 3652 | 52.60 | 23 |
| BB19 | 4.07 | 273 | 132 | 88 | 3778 | 41.30 | 13 |
| BB20 | 4.49 | 73 | 19 | 338 | 3725 | 37.90 | 13 |
| BB21 | 2.89 | 145 | 51 | 186 | 2812 | 44.40 | 8 |
| BB22 | 5.23 | 124 | 28 | 294 | 4210 | 34.20 | 32 |
| BB23 | 3.72 | 128 | 30 | 190 | 3427 | 23.40 | 9 |
| BB24 | 4.64 | 57 | 16 | 582 | 4438 | 31.30 | 58 |
| BB25 | 3.31 | 211 | 67 | 118 | 3069 | 44.30 | 8 |
| BB26 | 3.59 | 77 | 23 | 242 | 3547 | 44.60 | 8 |
| BB27 | 2.58 | 39 | 9 | 514 | 2602 | 43.10 | 9 |
| BB28 | 3.02 | 103 | 27 | 187 | 3056 | 26.30 | 4 |
| BB29 | 4.16 | 122 | 21 | 304 | 3375 | 34.40 | 21 |
| BB30 | 1.99 | 35 | 9 | 292 | 2109 | 25.80 | 4 |
| BB31 | 5.34 | 141 | 35 | 263 | 4342 | 45.30 | 15 |
| BB32 | 3.62 | 326 | 117 | 52 | 3061 | 35.20 | 14 |
| BB33 | 3.25 | 137 | 77 | 74 | 3319 | 56.80 | 9 |
| BB34 | 4.87 | 115 | 31 | 302 | 4339 | 38.20 | 9 |
| BB35 | 3.13 | 1.080 | 834 | 7 | 2773 | 31.80 | 12 |
| BB36 | 4.29 | 149 | 54 | 138 | 3907 | 39.10 | 9 |
| BB37 | 3.94 | 120 | 29 | 353 | 3831 | 51.50 | 11 |
| BB38 | 3.62 | 242 | 62 | 144 | 3260 | 44.20 | 14 |
| BB39 | 4.06 | 50 | 7 | 814 | 3832 | 59.10 | 17 |
| BB40 | 6.73 | 296 | 113 | 106 | 5886 | 45.10 | 0 |
| BB41 | 2.54 | 64 | 13 | 288 | 2631 | 32.90 | 7 |
| BB42 | 3.59 | 118 | 30 | 186 | 3352 | 33.10 | 23 |
| BB43 | 3.69 | 317 | 19 | 330 | 3767 | 49.10 | 10 |
| BB44 | 4.03 | 21 | 5 | 985 | 4077 | 40.20 | 17 |
| OB1 | 4.21 | 150 | 53 | 143 | 3949 | 43.10 | 13 |
| OB2 | 4.15 | 563 | 111 | 53 | 3804 | 47.20 | 14 |
| OB3 | 3.96 | 2.090 | 2.090 | 150 | 3665 | 48.70 | 16 |
| OB4 | 2.73 | 66.9 | 15 | 778 | 2962 | 38.90 | 4 |
| OB5 | 2.05 | 107 | 50 | 81 | 2113 | 31.90 | 5 |
| OB6 | 3.88 | 180 | 68 | 100 | 3643 | 39.10 | 11 |
| OB7 | 1.78 | 24 | 5 | 586 | 1916 | 26.10 | 2 |
| OB8 | 2.92 | 106 | 17 | 269 | 3159 | 31.60 | 7 |
| OB9 | 3.13 | 432 | 190 | 55 | 3181 | 55.10 | 6 |
| OB10 | 3 | 55 | 14 | 316 | 2910 | 48.60 | 10 |
| OB11 | 2.69 | 324 | 101 | 212 | 2676 | 30.50 | 5 |
| OB12 | 4.68 | 59 | 21 | 361 | 4128 | 39.90 | 22 |
| OB13 | 6.74 | 32 | 7 | 1266 | 5928 | 34.40 | 82 |
| OB14 | 6.81 | 436 | 121 | 365 | 6165 | 44.90 | 17 |
| OB15 | 2.25 | 20 | 4 | 800 | 2345 | 36.10 | 13 |

**Supplementary table S4:** *Colwellia* genomes used in this study for comparison to MAG BB5.

| Name | NCBI BioProject accession number | Isolation source | Number of M9 collagenases |
| --- | --- | --- | --- |
| *Colwellia piezophila* | PRJNA182419 | Deep-sea sediment | 2 |
| *Colwellia psychrerythraea* | PRJNA258170 | *Terua* mussel | 11 |
| *Colwellia hornerae* | PRJNA516280 | Arctic sea ice | 0 |
| *Colwellia demingiae* | PRJNA516284 | Arctic sea ice | 1 |
| Candidatus *Colwellia aromaticivorans* | PRJNA478776 | Microcosm experiments with oil in seawater | 0 |
| *Colwellia echini* | PRJNA420580 | Sea urchin | 0 |
| *Colwellia beringensis* | PRJNA378583 | Marine sediment, Bering Sea | 1 |
| *Colwellia agarivorans* | PRJNA371543 | Coastal sea water | 0 |
| *Colwellia marinimaniae* | PRJDB5767 | Amphipod from Challenger Deep | 4 |
| *Colwellia sediminilitoris* | PRJNA381102 | Tidal flat, South Sea, South Korea | 0 |
| *Colwellia polaris* | PRJNA380006 | Arctic sea ice | 0 |
| *Colwellia mytili* | PRJNA381102 | Mussel *Mytilus edulis* | 2 |
| *Colwellia aestuarii* | PRJNA371561 | Tidal flat Korea | 0 |
| *Colwellia chukchiensis* | PRJNA380006 | Arctic ocean | 1 |

### Supplementary figures

**Supplementary figure S1:** A) ROV image of bone incubation experiment in the Byfjorden at 68 m depth, shown is a cow tibia. B) Bones retrieved from the Byfjorden after nine months of incubation, bacterial mats and blackening at the epiphysis can be seen.

**Supplementary figure S2:** Maximum-likelihood tree of all 94 obtained carbonic anhydrases and relevant reference sequences from Capasso *et al.,* 2015 (40).

**Supplementary figure S3:** Genomic context of all identified M9 collagenase in the investigated MAGs. M9 collagenases are encircled in red, all genes potentially involved in collagen/proline utilization pathways are encircled in green. The graphic was made with SnapGene software (from GSL Biotech; available at snapgene.com).

**Supplementary figure S4:** Comparison of the bone microbiome to seawater metagenomes. Tara Oceans seawater MAGs have been chosen for comparison. The Tara Oceans dataset comprises here 832 bacterial MAGs with >70% completion. The bone microbiome dataset has been rebinned to >70% completion and contains now 86 MAGs for this comparison. A) Percental bacterial class abundance comparison between bone microbiome MAGs (orange) and Tara Oceans MAGs (blue). Only bacterial classes present within the bone microbiome have been included in this graph, all classes only present in the Tara Oceans dataset are combined in, Others’. B) Enzyme abundance comparison. Shown is the ratio per MAG for each investigated enzyme class. All MAGs have been profiled with the here established HMM profiles and the total obtained number of enzymes was divided by the number of MAGs per dataset. Highlighted in red are the higher ratios and in bold red ratios at least twice as high as in the other datatset.

## References

1. Debashish G, Malay S, Barindra S, Joydeep M. 2005. Marine enzymes. Adv Biochem Eng Biotechnol 96:189–218.

2. Imhoff JF, Labes A, Wiese J. 2011. Bio-mining the microbial treasures of the ocean: new natural products. Biotechnol Adv 29:468–82.

3. Trincone A. 2011. Marine biocatalysts: enzymatic features and applications. Mar Drugs 9:478–99.

4. Adams MW, Perler FB, Kelly RM. 1995. Extremozymes: expanding the limits of biocatalysis. Biotechnology (N Y) 13:662–8.

5. Aertsen A, Meersman F, Hendrickx ME, Vogel RF, Michiels CW. 2009. Biotechnology under high pressure: applications and implications. Trends Biotechnol 27:434–41.

6. Feller G, Gerday C. 2003. Psychrophilic enzymes: hot topics in cold adaptation. Nat Rev Microbiol 1:200–8.

7. Alcaide M, Tchigvintsev A, Martínez-Martínez M, Popovic A, Reva ON, Lafraya Á, Bargiela R, Nechitaylo TY, Matesanz R, Cambon-Bonavita MA, Jebbar M, Yakimov MM, Savchenko A, Golyshina OV, Yakunin AF, Golyshin PN, Ferrer M, Consortium M. 2015. Identification and characterization of carboxyl esterases of gill chamber-associated microbiota in the deep-sea shrimp *Rimicaris exoculata* by using functional metagenomics. Appl Environ Microbiol 81:2125–36.

8. Kennedy J, O’Leary ND, Kiran GS, Morrissey JP, O’Gara F, Selvin J, Dobson AD. 2011. Functional metagenomic strategies for the discovery of novel enzymes and biosurfactants with biotechnological applications from marine ecosystems. J Appl Microbiol 111:787–99.

9. Popovic A, Tchigvintsev A, Tran H, Chernikova TN, Golyshina OV, Yakimov MM, Golyshin PN, Yakunin AF. 2015. Metagenomics as a tool for enzyme discovery: hydrolytic enzymes from marine-related metagenomes. Adv Exp Med Biol 883:1–20.

10. Jørgensen BB, Boetius A. 2007. Feast and famine--microbial life in the deep-sea bed. Nat Rev Microbiol 5:770–81.

11. Naganuma T. 2000. Microbes on the edge of global biosphere. Biol Sci Space 14:323–31.

12. Sogin ML, Morrison HG, Huber JA, Mark Welch D, Huse SM, Neal PR, Arrieta JM, Herndl GJ. 2006. Microbial diversity in the deep sea and the underexplored “rare biosphere”. Proc Natl Acad Sci U S A 103:12115–20.

13. Smith C, Baco A. 2003. Ecology of whale falls at the deep-sea floor, p 311–354. In Gibson R, Atkinson R (ed), Oceanography and Marine Biology: an Annual Review, vol 41. Taylor&Francis.

14. Rouse GW, Goffredi SK, Vrijenhoek RC. 2004. *Osedax:* bone-eating marine worms with dwarf males. Science 305:668–71.

15. Miyamoto N, Yoshida MA, Koga H, Fujiwara Y. 2017. Genetic mechanisms of bone digestion and nutrient absorption in the bone-eating worm *Osedax japonicus* inferred from transcriptome and gene expression analyses. BMC Evol Biol 17:17.

16. Tresguerres M, Katz S, Rouse GW. 2013. How to get into bones: proton pump and carbonic anhydrase in *Osedax* boneworms. Proc Biol Sci 280:20130625.

17. Higgs ND, Glover AG, Dahlgren TG, Little CT. 2011. Bone-boring worms: characterizing the morphology, rate, and method of bioerosion by *Osedax mucofloris* (Annelida, Siboglinidae). Biol Bull 221:307–16.

18. Goffredi SK, Yi H, Zhang Q, Klann JE, Struve IA, Vrijenhoek RC, Brown CT. 2014. Genomic versatility and functional variation between two dominant heterotrophic symbionts of deep-sea *Osedax* worms. ISME J 8:908–24.

19. Goffredi SK, Orphan VJ, Rouse GW, Jahnke L, Embaye T, Turk K, Lee R, Vrijenhoek RC. 2005. Evolutionary innovation: a bone-eating marine symbiosis. Environ Microbiol 7:1369–78.

20. Goffredi SK, Johnson SB, Vrijenhoek RC. 2007. Genetic diversity and potential function of microbial symbionts associated with newly discovered species of *Osedax* polychaete worms. Appl Environ Microbiol 73:2314–23.

21. Verna C, Ramette A, Wiklund H, Dahlgren TG, Glover AG, Gaill F, Dubilier N. 2010. High symbiont diversity in the bone-eating worm *Osedax mucofloris* from shallow whale-falls in the North Atlantic. Environ Microbiol 12:2355–70.

22. Glover AG, Källström B, Smith CR, Dahlgren TG. 2005. World-wide whale worms? A new species of *Osedax* from the shallow north Atlantic. Proc Biol Sci 272:2587–92.

23. Dahlgren T, Wiklund H, Källström B, Lundälv T, Smith C, Glover A. 2006. A shallowwater whale-fall experiment in the North Atlantic. Cahiers De Biologie Marine 47:385–389.

24. Robey PG. 2008. Noncollagenous bone matrix proteins, p 335–349. In Bilezikian JP, Raisz LG, Martin JT (ed), Principles of bone biology, Third edition ed, vol Volume I. Academic press.

25. Sato S, Rahemtulla F, Prince CW, Tomana M, Butler WT. 2009. Acidic glycoproteins from bovine compact bone. Connective Tissue Research 14:51–64.

26. Kuboki Y, Watanabe T, Tazaki M, Taktta H. 1991. Comparative biochemistry of bone matrix proteins in bovine and fish. In Suga S, Nakahara H (ed), Mechanisms and phylogeny of mineralization in biological systems doi:https://doi.org/10.1007/978-4-431-68132-8_79. Springer, Tokyo.

27. Shoulders MD, Raines RT. 2009. Collagen structure and stability. Annu Rev Biochem 78:929–58.

28. de Gonzalo G, Colpa DI, Habib MH, Fraaije MW. 2016. Bacterial enzymes involved in lignin degradation. J Biotechnol 236:110–9.

29. Puentes-Tellez PE, Salles JF. 2018. Construction of effective minimal active microbial consortia for lignocellulose degradation. Microbial Ecology 76:419–429.

30. Vietti L, Bailey J, Ricci E. 2014. Insights into the microbial degradation of bone in marine environments from rRNA gene sequencing of biofilms on lab-simulated carcass-falls. The Paleontological Society Special Publications doi:doi:10.1017/S2475262200012582:120-121.

31. Vietti LA. 2014. Insights into the microbial degradation of bones from the marine vertebrate fossil record: an experimental approach using interdisciplinary analyses. Ph.D. dissertation. University of Minnesota, University of Minnesota Digital Conservancy.

32. Taboada S, Bas M, Avila C, Riesgo A. 2020. Phylogenetic characterization of marine microbial biofilms associated with mammal bones in temperate and polar areas. Marine Biodiversity 50.

33. Hewitt OH, Diez-Vives C, Taboada S. 2020. Microbial insights from Antarctic and Mediterranean shallow-water bone-eating worms. Polar Biology 43:1605–1621.

34. Freitas RC, Marques HIF, Silva MACD, Cavalett A, Odisi EJ, Silva BLD, Montemor JE, Toyofuku T, Kato C, Fujikura K, Kitazato H, Lima AOS. 2019. Evidence of selective pressure in whale fall microbiome proteins and its potential application to industry. Mar Genomics 45:21–27.

35. Menzel P, Ng KL, Krogh A. 2016. Fast and sensitive taxonomic classification for metagenomics with Kaiju. Nature Communications 7.

36. Parks DH, Chuvochina M, Waite DW, Rinke C, Skarshewski A, Chaumeil PA, Hugenholtz P. 2018. A standardized bacterial taxonomy based on genome phylogeny substantially revises the tree of life. Nat Biotechnol 36:996–1004.

37. De Anda V, Zapata-Peñasco I, Poot-Hernandez AC, Eguiarte LE, Contreras-Moreira B, Souza V. 2017. MEBS, a software platform to evaluate large (meta)genomic collections according to their metabolic machinery: unraveling the sulfur cycle. Gigascience 6:1–17.

38. Waite DW, Chuvochina M, Pelikan C, Parks DH, Yilmaz P, Wagner M, Loy A, Naganuma T, Nakai R, Whitman WB, Hahn MW, Kuever J, Hugenholtz P. 2020. Proposal to reclassify the proteobacterial classes Deltaproteobacteria and Oligoflexia, and the phylum Thermodesulfobacteria into four phyla reflecting major functional capabilities. Int J Syst Evol Microbiol doi:10.1099/ijsem.0.004213.

39. Müller AL, Kjeldsen KU, Rattei T, Pester M, Loy A. 2015. Phylogenetic and environmental diversity of DsrAB-type dissimilatory (bi)sulfite reductases. ISME J 9:1152–65.

40. Weimann A, Mooren K, Frank J, Pope PB, Bremges A, McHardy AC. 2016. From genomes to phenotypes: Traitar, the Microbial Trait Analyzer. mSystems 1.

41. Yamamoto M, Takai K. 2011. Sulfur metabolisms in epsilon- and gamma-proteobacteria in deep-sea hydrothermal fields. Front Microbiol 2:192.

42. Muyzer G, Stams AJ. 2008. The ecology and biotechnology of sulphate-reducing bacteria. Nat Rev Microbiol 6:441–54.

43. Barton LL, Fauque GD. 2009. Biochemistry, physiology and biotechnology of sulfatereducing bacteria. Adv Appl Microbiol 68:41–98.

44. Kruse S, Goris T, Westermann M, Adrian L, Diekert G. 2018. Hydrogen production by *Sulfurospirillum* species enables syntrophic interactions of Epsilonproteobacteria. Nat Commun 9:4872.

45. Schulz HN, Jorgensen BB. 2001. Big bacteria. Annu Rev Microbiol 55:105–37.

46. Gevertz D, Telang AJ, Voordouw G, Jenneman GE. 2000. Isolation and characterization of strains CVO and FWKO B, two novel nitrate-reducing, sulfide-oxidizing bacteria isolated from oil field brine. Appl Environ Microbiol 66:2491–501.

47. Trojan D, Schreiber L, Bjerg JT, Bøggild A, Yang T, Kjeldsen KU, Schramm A. 2016. A taxonomic framework for cable bacteria and proposal of the candidate genera *Electrothrix* and *Electronema*. Syst Appl Microbiol 39:297–306.

48. Pfeffer C, Larsen S, Song J, Dong M, Besenbacher F, Meyer RL, Kjeldsen KU, Schreiber L, Gorby YA, El-Naggar MY, Leung KM, Schramm A, Risgaard-Petersen N, Nielsen LP. 2012. Filamentous bacteria transport electrons over centimetre distances. Nature 491:218–21.

49. Capasso C, Supuran CT. 2015. An overview of the alpha-, beta- and gamma-carbonic anhydrases from Bacteria: can bacterial carbonic anhydrases shed new light on evolution of bacteria? J Enzyme Inhib Med Chem 30:325–32.

50. Bowman JP, Gosnik JJ, McCammon SA, Lewis TE, Nichols DS, Nichols PD, Skerratt H, Staley JT, McMeekin TA. 1998. *Colwellia demingiae* sp. nov., *Colwellia hornerae* sp. nov., *Colwellia rossensis* sp. nov. and *Colwellia psychrotropica* sp. nov.: psychrophilic Antarctic species with the ability to synthesize docosahexaenoic acid (22:ω63). International Journal of Systematic and Evolutionary Microbiology 48:1171–1180.

51. Bowman JP, McCammon SA, Brown MV, Nichols DS, McMeekin TA. 1997. Diversity and association of psychrophilic bacteria in Antarctic sea ice. Appl Environ Microbiol 63:3068–78.

52. Campeao ME, Swings J, Silva BS, Otsuki K, Thompson FL, Thompson CC. 2019. *“Candidatus Colwellia aromaticivorans”* sp. nov., *“Candidatus Halocyntiibacter alkanivorans”* sp. nov., and *“Candidatus Ulvibacter alkanivorans”* sp. nov. Genome Sequences. Microbiol Resour Announc 8.

53. Christiansen L, Bech PK, Schultz-Johansen M, Martens HJ, Stougaard P. 2018. *Colwellia echini* sp. nov., an agar- and carrageenan-solubilizing bacterium isolated from sea urchin. Int J Syst Evol Microbiol 68:687–691.

54. Jung SY, Oh TK, Yoon JH. 2006. *Colwellia aestuarii* sp. nov., isolated from a tidal flat sediment in Korea. Int J Syst Evol Microbiol 56:33–7.

55. Kim YO, Park IS, Park S, Nam BH, Jung YT, Kim DG, Yoon JH. 2017. *Colwellia mytili* sp. nov., isolated from mussel *Mytilus edulis*. Int J Syst Evol Microbiol 67:31–36.

56. Kusube M, Kyaw TS, Tanikawa K, Chastain RA, Hardy KM, Cameron J, Bartlett DH. 2017. *Colwellia marinimaniae* sp. nov., a hyperpiezophilic species isolated from an amphipod within the Challenger Deep, Mariana Trench. Int J Syst Evol Microbiol 67:824–831.

57. Nogi Y, Hosoya S, Kato C, Horikoshi K. 2004. *Colwellia piezophila* sp. nov., a novel piezophilic species from deep-sea sediments of the Japan Trench. Int J Syst Evol Microbiol 54:1627–1631.

58. Park S, Jung YT, Yoon JH. 2016. *Colwellia sediminilitoris* sp. nov., isolated from a tidal flat. Int J Syst Evol Microbiol 66:3258–3263.

59. Techtmann SM, Fitzgerald KS, Stelling SC, Joyner DC, Uttukar SM, Harris AP, Alshibli NK, Brown SD, Hazen TC. 2016. *Colwellia psychrerythraea* strains from distant deep sea basins show adaptation to local conditions. Frontiers in Environmental Science 4.

60. Xu ZX, Zhang HX, Han JR, Dunlap CA, Rooney AP, Mu DS, Du ZJ. 2017. *Colwellia agarivorans* sp. nov., an agar-digesting marine bacterium isolated from coastal seawater. Int J Syst Evol Microbiol 67:1969–1974.

61. Yu Y, Li HR, Zeng YX. 2011. *Colwellia chukchiensis* sp. nov., a psychrotolerant bacterium isolated from the Arctic Ocean. Int J Syst Evol Microbiol 61:850–853.

62. Zhang C, Guo W, Wang Y, Chen X. 2017. *Colwellia beringensis* sp. nov., a psychrophilic bacterium isolated from the Bering Sea. Int J Syst Evol Microbiol 67:5102–5107.

63. Zhang DC, Yu Y, Xin YH, Liu HC, Zhou PJ, Zhou YG. 2008. *Colwellia polaris* sp. nov., a psychrotolerant bacterium isolated from Arctic sea ice. Int J Syst Evol Microbiol 58:1931–4.

64. Delmont TO, Quince C, Shaiber A, Esen OC, Lee STM, Rappe MS, McLellan SL, Lucker S, Eren AM. 2018. Nitrogen-fixing populations of Planctomycetes and Proteobacteria are abundant in surface ocean metagenomes (vol 3, pg 804, 2018). Nature Microbiology 3.

65. Gontang EA, Fenical W, Jensen PR. 2007. Phylogenetic diversity of gram-positive bacteria cultured from marine sediments. Appl Environ Microbiol 73:3272–82.

66. Teske A, Brinkhoff T, Muyzer G, Moser DP, Rethmeier J, Jannasch HW. 2000. Diversity of thiosulfate-oxidizing bacteria from marine sediments and hydrothermal vents. Appl Environ Microbiol 66:3125–33.

67. Leloup J, Fossing H, Kohls K, Holmkvist L, Borowski C, Jørgensen BB. 2009. Sulfatereducing bacteria in marine sediment (Aarhus Bay, Denmark): abundance and diversity related to geochemical zonation. Environ Microbiol 11:1278–91.

68. Cotter PD, Hill C. 2003. Surviving the acid test: responses of gram-positive bacteria to low pH. Microbiol Mol Biol Rev 67:429–53, table of contents.

69. Anné J, Economou A, Bernaerts K. 2017. Protein secretion in gram-positive bacteria: From multiple pathways to biotechnology. Curr Top Microbiol Immunol 404:267–308.

70. Jang H, Yang SH, Seo HS, Lee JH, Kim SJ, Kwon KK. 2015. *Amphritea spongicola* sp nov., isolated from a marine sponge, and emended description of the genus *Amphritea*. International Journal of Systematic and Evolutionary Microbiology 65:1866–1870.

71. Wang JH, Lu Y, Nawaz MZ, Xu J. 2018. Comparative genomics reveals evidence of genome reduction and high extracellular protein degradation potential in *Kangiella*. Frontiers in Microbiology 9.

72. Cao SN, He JF, Zhang F, Lin L, Gao Y, Zhou QM. 2019. Diversity and community structure of bacterioplankton in surface waters off the northern tip of the Antarctic Peninsula. Polar Research 38.

73. Lastovica A, On S, Zhang L. 2014. The family *Campylobacteraceae*, p 307–335. *In* Rosenburg E, DeLong E, Lory S, Stackebrandt E, Thompson F (ed), The Prokaryotes doi:https://doi.org/10.1007/978-3-642-39044-9_274. Springer.

74. Takai K, Campbell BJ, Cary SC, Suzuki M, Oida H, Nunoura T, Hirayama H, Nakagawa S, Suzuki Y, Inagaki F, Horikoshi K. 2005. Enzymatic and genetic characterization of carbon and energy metabolisms by deep-sea hydrothermal chemolithoautotrophic isolates of Epsilonproteobacteria. Appl Environ Microbiol 71:7310–20.

75. Deming JW, Reysenbach AL, Macko SA, Smith CR. 1997. Evidence for the microbial basis of a chemoautotrophic invertebrate community at a whale fall on the deep seafloor: bone-colonizing bacteria and invertebrate endosymbionts. Microsc Res Tech 37:162–70.

76. Supuran CT, Capasso C. 2017. An overview of the bacterial carbonic anhydrases. Metabolites 7.

77. Vietti L, Bailey J, Fox D, Rogers R. 2015. Rapid formation of framboidal sulfides on bone surfaces from a simulated marine carcass fall. PALAIOS 30:327–334.

78. Naretto A, Fanuel M, Ropartz D, Rogniaux H, Larocque R, Czjzek M, Tellier C, Michel G. 2019. The agar-specific hydrolase. J Biol Chem 294:6923–6939.

79. Nedashkovskaya OI, Kim SG, Stenkova AM, Kukhlevskiy AD, Zhukova NV, Mikhailov VV. 2018. *Aquimarina algiphila* sp. nov., a chitin degrading bacterium isolated from the red alga *Tichocarpus crinitus*. Int J Syst Evol Microbiol 68:892–898.

80. Konasani VR, Jin C, Karlsson NG, Albers E. 2018. A novel ulvan lyase family with broad-spectrum activity from the ulvan utilisation loci of *Formosa agariphila* KMM 3901. Sci Rep 8:14713.

81. Li S, Wang L, Chen X, Zhao W, Sun M, Han Y. 2018. Cloning, expression, and biochemical characterization of two new oligoalginate lyases with synergistic degradation capability. Mar Biotechnol (NY) 20:75–86.

82. Shen J, Chang Y, Chen F, Dong S. 2018. Expression and characterization of a κ-carrageenase from marine bacterium *Wenyingzhuangia aestuarii* OF219: A biotechnological tool for the depolymerization of κ-carrageenan. Int J Biol Macromol 112:93–100.

83. Tan H, Miao R, Liu T, Yang L, Yang Y, Chen C, Lei J, Li Y, He J, Sun Q, Peng W, Gan B, Huang Z. 2018. A bifunctional cellulase-xylanase of a new *Chryseobacterium* strain isolated from the dung of a straw-fed cattle. Microb Biotechnol 11:381–398.

84. Rochat T, Pérez-Pascual D, Nilsen H, Carpentier M, Bridel S, Bernardet JF, Duchaud E. 2019. Identification of a novel elastin-degrading enzyme from the fish pathogen. Appl Environ Microbiol 85.

85. Choi KD, Lee GE, Park JS. 2018. *Aquimarina spongiicola* sp. nov., isolated from spongin. Int J Syst Evol Microbiol 68:990–994.

86. Iino T, Mori K, Itoh T, Kudo T, Suzuki K, Ohkuma M. 2014. Description of *Mariniphaga anaerophila* gen. nov., sp. nov., a facultatively aerobic marine bacterium isolated from tidal flat sediment, reclassification of the Draconibacteriaceae as a later heterotypic synonym of the Prolixibacteraceae and description of the family Marinifilaceae fam. nov. Int J Syst Evol Microbiol 64:3660–7.

87. Han Z, Shang-Guan F, Yang J. 2019. Molecular and biochemical characterization of a bimodular xylanase From. Front Microbiol 10:1507.

88. Emmons A, Mundorff A, Keenan S, Davoren J, Andronowski J, Carter D, DeBruyn J. 2019. Patterns of microbial colonization of human bone from surface-decomposed remains. bioRxiv 664482.

89. Robey P. 2002. Bone matrix proteoglycans and glycoproteins. *In* Bilezikian J, Raisz L, Rodan G (ed), Principles of bone biology, 2nd Edition ed doi:https://doi.org/10.1016/B978-012098652-1.50116-5. Elsevier.

90. van Vliet DM, Palakawong Na Ayudthaya S, Diop S, Villanueva L, Stams AJM, Sánchez-Andrea I. 2019. Anaerobic degradation of sulfated polysaccharides by two novel. Front Microbiol 10:253.

91. S D, O G. 2016. *Colwellia* and sulfur-oxidizing bacteria: An unusual dual symbiosis in a *Terua* mussel (Mytilidae: Bathymodiolinae) from whale falls in the Antilles arc. Deep Sea Research Part I 115:112–122.

92. Hansen AM, Qiu Y, Yeh N, Blattner FR, Durfee T, Jin DJ. 2005. SspA is required for acid resistance in stationary phase by downregulation of H-NS in *Escherichia coli*. Mol Microbiol 56:719–34.

93. Bolger AM, Lohse M, Usadel B. 2014. Trimmomatic: a flexible trimmer for Illumina sequence data. Bioinformatics 30:2114–20.

94. Bankevich A, Nurk S, Antipov D, Gurevich AA, Dvorkin M, Kulikov AS, Lesin VM, Nikolenko SI, Pham S, Prjibelski AD, Pyshkin AV, Sirotkin AV, Vyahhi N, Tesler G, Alekseyev MA, Pevzner PA. 2012. SPAdes: a new genome assembly algorithm and its applications to single-cell sequencing. J Comput Biol 19:455–77.

95. Uritskiy GV, DiRuggiero J, Taylor J. 2018. MetaWRAP-a flexible pipeline for genome-resolved metagenomic data analysis. Microbiome 6:158.

96. Alneberg J, Bjarnason BS, de Bruijn I, Schirmer M, Quick J, Ijaz UZ, Lahti L, Loman NJ, Andersson AF, Quince C. 2014. Binning metagenomic contigs by coverage and composition. Nat Methods 11:1144–6.

97. Wu YW, Simmons BA, Singer SW. 2016. MaxBin 2.0: an automated binning algorithm to recover genomes from multiple metagenomic datasets. Bioinformatics 32:605–7.

98. Kang DD, Li F, Kirton E, Thomas A, Egan R, An H, Wang Z. 2019. MetaBAT 2: an adaptive binning algorithm for robust and efficient genome reconstruction from metagenome assemblies. PeerJ 7:e7359.

99. Parks DH, Imelfort M, Skennerton CT, Hugenholtz P, Tyson GW. 2015. CheckM: assessing the quality of microbial genomes recovered from isolates, single cells, and metagenomes. Genome Res 25:1043–55.

100. Hyatt D, Chen GL, Locascio PF, Land ML, Larimer FW, Hauser LJ. 2010. Prodigal: prokaryotic gene recognition and translation initiation site identification. BMC Bioinformatics 11:119.

101. Huerta-Cepas J, Forslund K, Coelho LP, Szklarczyk D, Jensen LJ, von Mering C, Bork P. 2017. Fast genome-wide functional annotation through orthology assignment by eggNOG-Mapper. Mol Biol Evol 34:2115–2122.

102. Huerta-Cepas J, Szklarczyk D, Forslund K, Cook H, Heller D, Walter MC, Rattei T, Mende DR, Sunagawa S, Kuhn M, Jensen LJ, von Mering C, Bork P. 2016. eggNOG 4.5: a hierarchical orthology framework with improved functional annotations for eukaryotic, prokaryotic and viral sequences. Nucleic Acids Res 44:D286–93.

103. Aziz RK, Bartels D, Best AA, DeJongh M, Disz T, Edwards RA, Formsma K, Gerdes S, Glass EM, Kubal M, Meyer F, Olsen GJ, Olson R, Osterman AL, Overbeek RA, McNeil LK, Paarmann D, Paczian T, Parrello B, Pusch GD, Reich C, Stevens R, Vassieva O, Vonstein V, Wilke A, Zagnitko O. 2008. The RAST Server: rapid annotations using subsystems technology. BMC Genomics 9:75.

104. Overbeek R, Olson R, Pusch GD, Olsen GJ, Davis JJ, Disz T, Edwards RA, Gerdes S, Parrello B, Shukla M, Vonstein V, Wattam AR, Xia F, Stevens R. 2014. The SEED and the Rapid Annotation of microbial genomes using Subsystems Technology (RAST). Nucleic Acids Res 42:D206–14.

105. Quevillon E, Silventoinen V, Pillai S, Harte N, Mulder N, Apweiler R, Lopez R. 2005. InterProScan: protein domains identifier. Nucleic Acids Research 33:W116–W120.

106. Caspi R, Billington R, Ferrer L, Foerster H, Fulcher CA, Keseler IM, Kothari A, Krummenacker M, Latendresse M, Mueller LA, Ong Q, Paley S, Subhraveti P, Weaver DS, Karp PD. 2016. The MetaCyc database of metabolic pathways and enzymes and the BioCyc collection of pathway/genome databases. Nucleic Acids Research 44:D471–D480.

107. Price MN, Dehal PS, Arkin AP. 2010. FastTree 2--approximately maximum-likelihood trees for large alignments. PLoS One 5:e9490.

108. Letunic I, Bork P. 2007. Interactive Tree Of Life (iTOL): an online tool for phylogenetic tree display and annotation. Bioinformatics 23:127–8.

109. Letunic I, Bork P. 2019. Interactive Tree Of Life (iTOL) v4: recent updates and new developments. Nucleic Acids Res 47:W256–W259.

110. Babicki S, Arndt D, Marcu A, Liang Y, Grant JR, Maciejewski A, Wishart DS. 2016. Heatmapper: web-enabled heat mapping for all. Nucleic Acids Res 44:W147–53.

111. Harrison KJ, Crécy-Lagard V, Zallot R. 2018. Gene Graphics: a genomic neighborhood data visualization web application. Bioinformatics 34:1406–1408.

112. Almagro Armenteros JJ, Tsirigos KD, Sønderby CK, Petersen TN, Winther O, Brunak S, von Heijne G, Nielsen H. 2019. SignalP 5.0 improves signal peptide predictions using deep neural networks. Nat Biotechnol 37:420–423.

113. Eddy SR. 2011. Accelerated Profile HMM Searches. PLoS Comput Biol 7:e1002195.

114. Larkin MA, Blackshields G, Brown NP, Chenna R, McGettigan PA, McWilliam H, Valentin F, Wallace IM, Wilm A, Lopez R, Thompson JD, Gibson TJ, Higgins DG. 2007. Clustal W and Clustal X version 2.0. Bioinformatics 23:2947–8.

115. Delmont TO, Quince C, Shaiber A, Esen OC, Lee STM, Rappe MS, MacLellan SL, Lucker S, Eren AM. 2018. Nitrogen-fixing populations of Planctomycetes and Proteobacteria are abundant in surface ocean metagenomes. Nature Microbiology 3:804–+.

