## Supplementary figures 1-4 for "Deciphering a marine bone degrading microbiome reveals a complex community effort"

**Supplementary figures for ‚Deciphering a marine bone degrading microbiome reveals a complex community effort”**

A

B

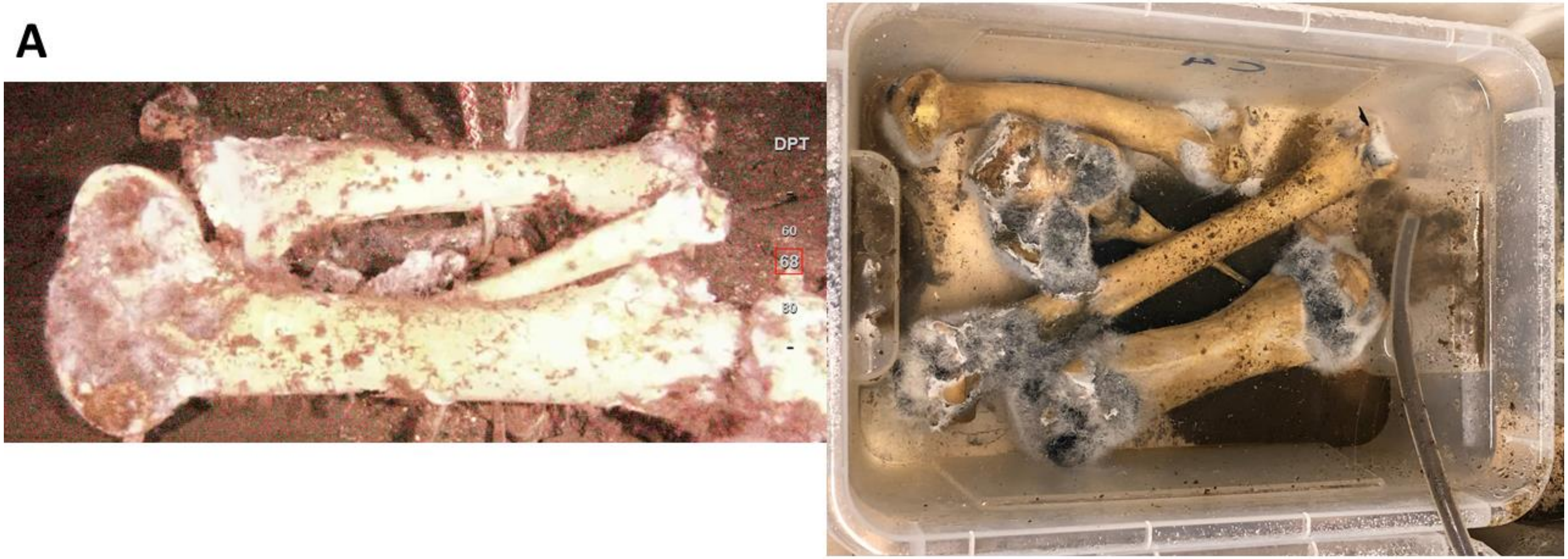


**Supplementary figure S1:** A) ROV image of bone incubation experiment in the Byfjorden at 68 m depth, shown is a cow tibia. B) Bones retrieved from the Byfjorden after nine months of incubation, bacterial mats and blackening at the epiphysis can be seen.

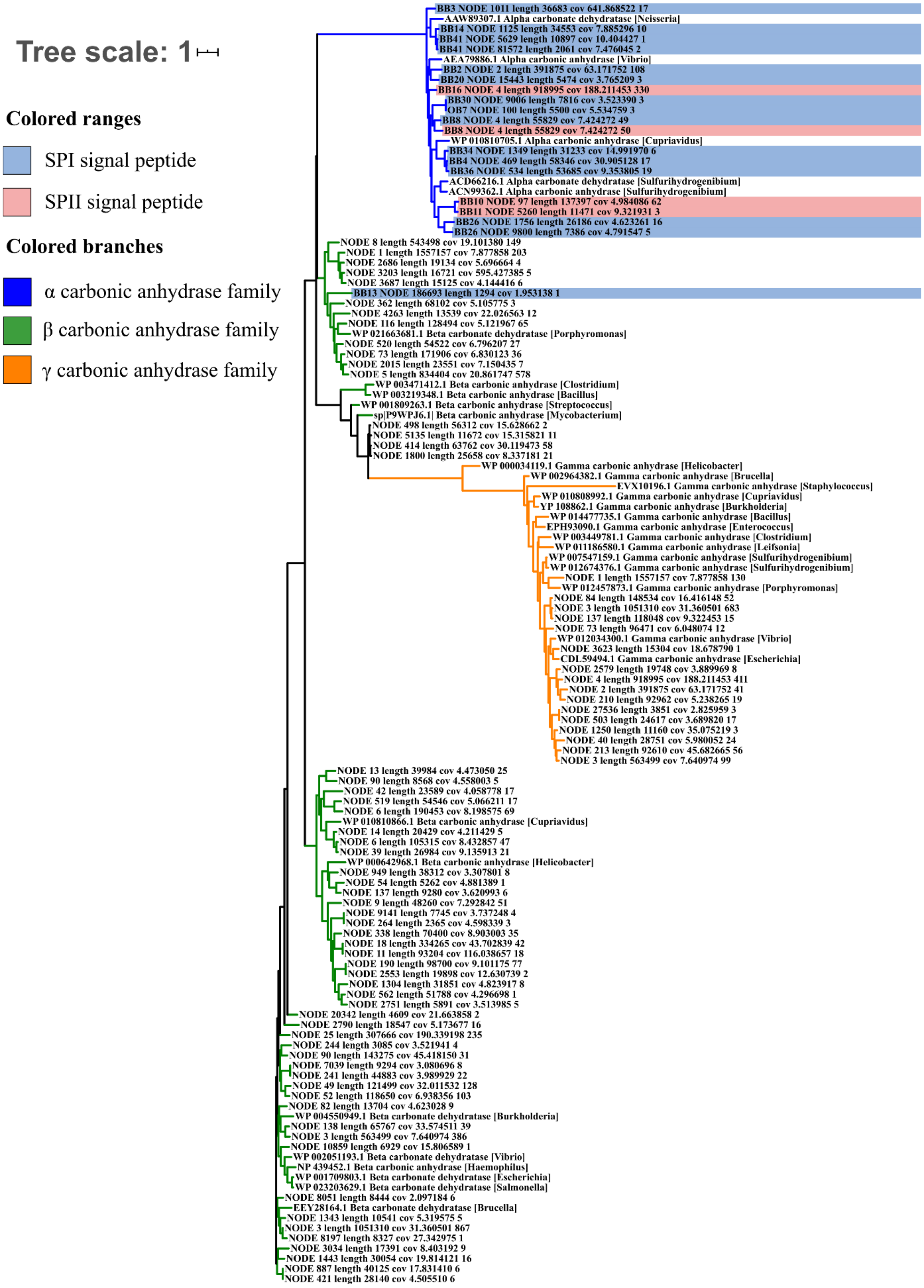


**Supplementary figure S2:** Maximum-likelihood tree of all 94 obtained carbonic anhydrases and relevant reference sequences from Capasso *et al.*, 2015 (40).

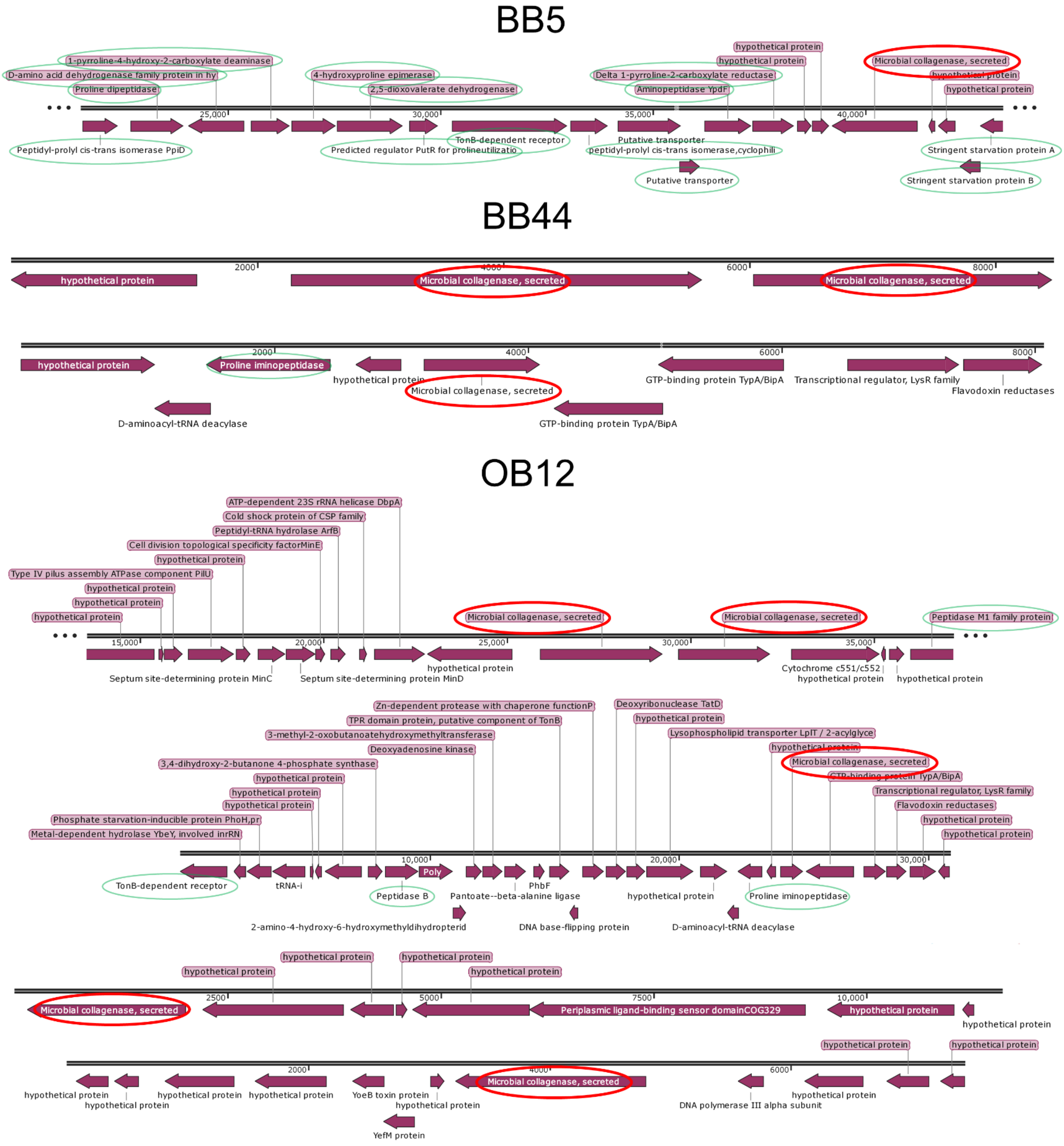


**Supplementary figure S3:** Genomic context of all identified M9 collagenase in the investigated MAGs. M9 collagenases are encircled in red, all genes potentially involved in collagen/proline utilization pathways are encircled in green. The graphic was made with SnapGene software (from GSL Biotech; available at snapgene.com).

**A**

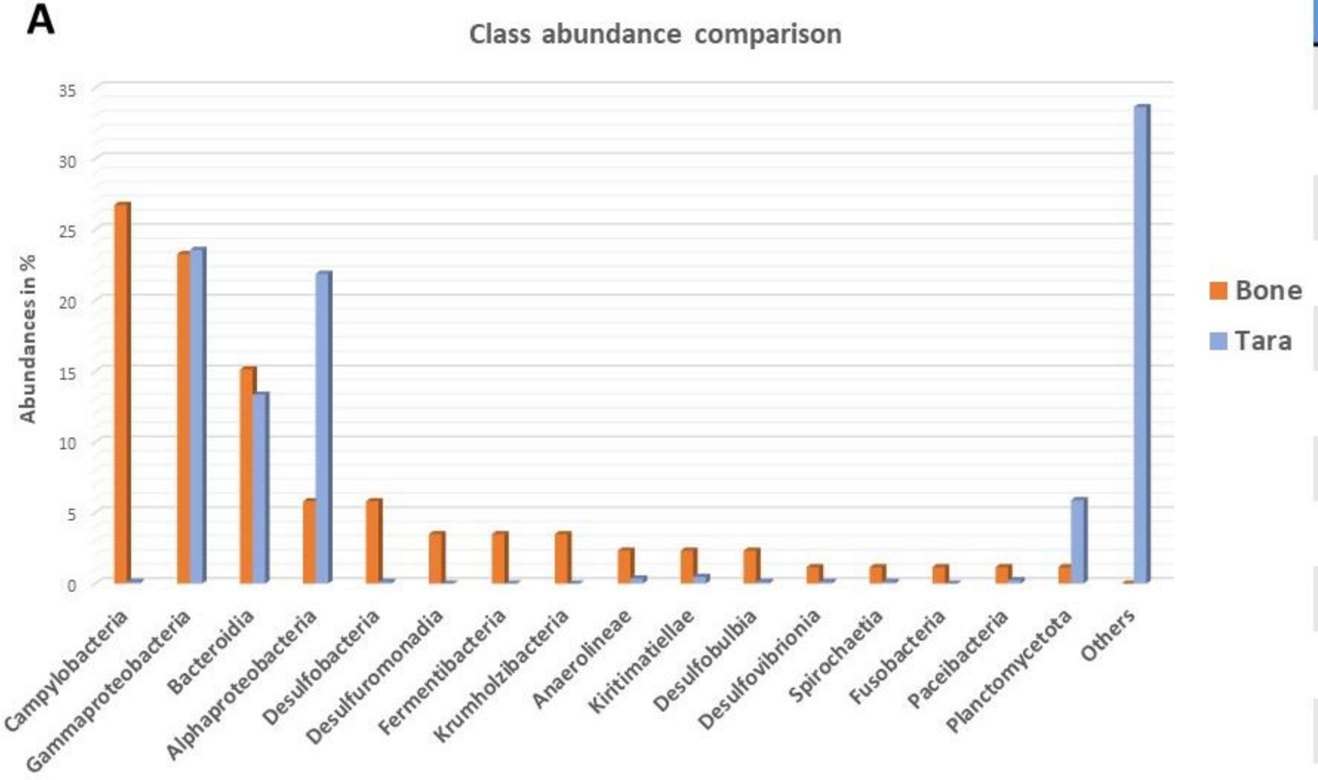


**B**

| Enzyme family | Ratio per MAG (Tara) | Ratio per MAG (Bone) |
|---|---|---|
| COG0383 | 0.09 | **0.19** |
| COG0673 | **5.34** | 1.67 |
| COG1472 | 1.05 | 1.15 |
| COG2303 | 1.6 | 0.94 |
| COG3250 | 0.45 | **1.03** |
| COG3669 | 0.21 | **0.56** |
| COG4409 | 0.24 | 0.26 |
| M9_Col | 0.002 | **0.13** |
| S1_Pep. | 2.03 | 2.03 |
| S8/S53_Pep. | 0.66 | 1 |
| U32_Pep. | 0.19 | **1.53** |
| PF16499 | 0.02 | **0.05** |

**Supplementary figure S4:** Comparison of the bone microbiome to seawater metagenomes. Tara Oceans seawater MAGs have been chosen for comparison. The Tara Oceans dataset comprises here 832 bacterial MAGs with >70% completion. The bone microbiome dataset has been rebinned to >70% completion and contains now 86 MAGs for this comparison. A) Percental bacterial class abundance comparison between bone microbiome MAGs (orange) and Tara Oceans MAGs (blue). Only bacterial classes present within the bone microbiome have been included in this graph, all classes only present in the Tara Oceans dataset are combined in ‚Others‘. B) Enzyme abundance comparison. Shown is the ratio per MAG for each investigated enzyme class. All MAGs have been profiled with the here established HMM profiles and the total obtained number of enzymes was divided by the number of MAGs per dataset. Highlighted in red are the higher ratios and in bold red ratios at least twice as high as in the other datatset.
